# A hierarchical framework for cortical and subcortical gray-matter parcellation across rodents, primates, and humans

**DOI:** 10.1101/2025.09.08.675002

**Authors:** Siva Venkadesh, Yuhe Tian, Wen-Jieh Linn, Jessica Barrios Martinez, Harrison Mansour, James Cook, David J. Schaeffer, Diego Szczupak, Afonso C. Silva, G Allan Johnson, Fang-Cheng Yeh

## Abstract

Translational neuroscience requires consistent anatomical frameworks to compare brain organization across species despite differences in size and specialization. Existing atlases are species-specific, limiting cross-species analyses. Here we developed a hierarchical common atlas delineating homologous cortical and subcortical gray matter regions across mouse, rat, marmoset, rhesus macaque, and human, built upon population-averaged minimal deformation templates and uniform tissue segmentation. The atlas was validated through cross-atlas Dice similarity against established human parcellations, cross-scale containment against species-specific atlases, cross-species geometric consistency of regional positioning, and comparison of independent mouse and marmoset invasive tracer connectivity, with a per-region homology confidence index quantifying the strength of each regional assignment. Cross-species tracer comparison revealed a structured gradient of correspondence, with sensorimotor connections showing strong conservation and association connections showing progressive divergence. This freely available atlas provides a unified coordinate system for comparative neuroscience, enabling quantitative evaluation of where cross-species correspondence holds and where species-specific divergence emerges.

## 1. Introduction

Translational neuroscience aims to uncover principles of brain organization that are shared across species, as well as those that have evolved for specific lineages. However, achieving this requires consistent definitions of brain regions across different species—a task complicated by substantial differences in brain size, folding, and specialization. Despite rapid advances in neuroimaging and histological mapping, a major obstacle persists: there is a need for standardized anatomical framework for translating brains across species. Most available atlases are species-specific and have been developed independently, often using different criteria for parcellation^1–4^, This leads to substantial differences in how homologous regions are defined, which can hinder our ability to determine whether observed similarities or differences in network organization reflect true biological principles or are simply artifacts of methodological variation.

Several initiatives have begun to address this challenge. For example, harmonized human atlases have been developed to reconcile differences across parcellation schemes^4–6^, and recent efforts have aligned nonhuman primate parcellations to human cortical surfaces to facilitate cross-species comparisons^7,8^. While these approaches have improved anatomical consistency within or between primates, comparable frameworks that encompass both rodents and primates are still lacking. Furthermore, most existing solutions are based on species-specific surfaces or templates and do not utilize population-averaged minimal deformation templates (MDTs), which are reference spaces created by iteratively averaging multiple individuals to minimize anatomical variability. In humans, the ICBM 2009a templates^9^ represent the most widely adopted standard, constructed through population averaging and nonlinear registration to reduce inter-individual anatomical differences. In contrast, atlases for other species often rely on single-subject scans or templates not generated using comparable MDT frameworks, resulting in inconsistencies in spatial alignment and anatomical scaling. This methodological disparity undermines the reliability of comparisons across species.

To address this gap, we first constructed population-averaged MDTs for each species to provide a consistent anatomical foundation for cross-species alignment. For humans and marmosets, we adopted existing MDTs from the widely used ICBM2009a templates^9^ and the Marmoset Brain Atlas^10^, respectively. For mouse, rat, and rhesus macaque, we generated new MDTs using high-resolution MRI and iterative diffeomorphic averaging (Online Methods). Building on these templates, we then delineated a common hierarchical atlas (CHA) of cortical and subcortical gray matter spanning rodents, nonhuman primates, and humans. Beyond the parcellation itself, we developed a quantitative validation framework including cross-atlas benchmarking, cross-scale containment analysis, cross-species geometric consistency, and independent tracer connectivity comparison, culminating in a per-region homology confidence index that enables transparent evaluation of each regional assignment.

The parcellation framework is organized hierarchically, beginning with an initial Level-0 tissue segmentation (defining major tissue compartments), followed by Level-1 macro-regions (major cortical and subcortical divisions), and Level-2 subdivisions (finer anatomical parcels), with all levels defined by consistent anatomical boundaries and literature-guided homologies. For closely related species (e.g., mouse and rat), parcellations were aligned using multimodal nonlinear spatial registration (see Online Methods), while for more divergent species (e.g., humans and nonhuman primates), boundaries were matched using anatomical landmarks and literature-based homologies. Validation confirmed robust regional correspondence through multiple independent approaches, including cross-atlas Dice similarity against established human parcellations, cross-scale containment against species-specific atlases, geometric consistency of regional positioning across species, and cross-species comparison of mouse and marmoset invasive tracer connectivity. We have made this atlas freely available to support comparative connectomics and translational studies, providing a unified coordinate system for formal cross-species evaluation of brain connectivity and organization.

## 2. Results

### 2.1. Overview of the Common Atlas Framework

Figure 1 summarizes the overall workflow for constructing the cross-species common atlas. Input MRI data from human, rhesus macaque, marmoset, rat and mouse were first used to generate Minimal Deformation Templates (MDTs) through iterative diffeomorphic averaging, similar to ANTs population-template methods^11,12^. The MDT ensures that each species’ reference template represents a population-average configuration with minimal anatomical distortion, providing a consistent spatial foundation for inter-species alignment.

**Figure 1.**
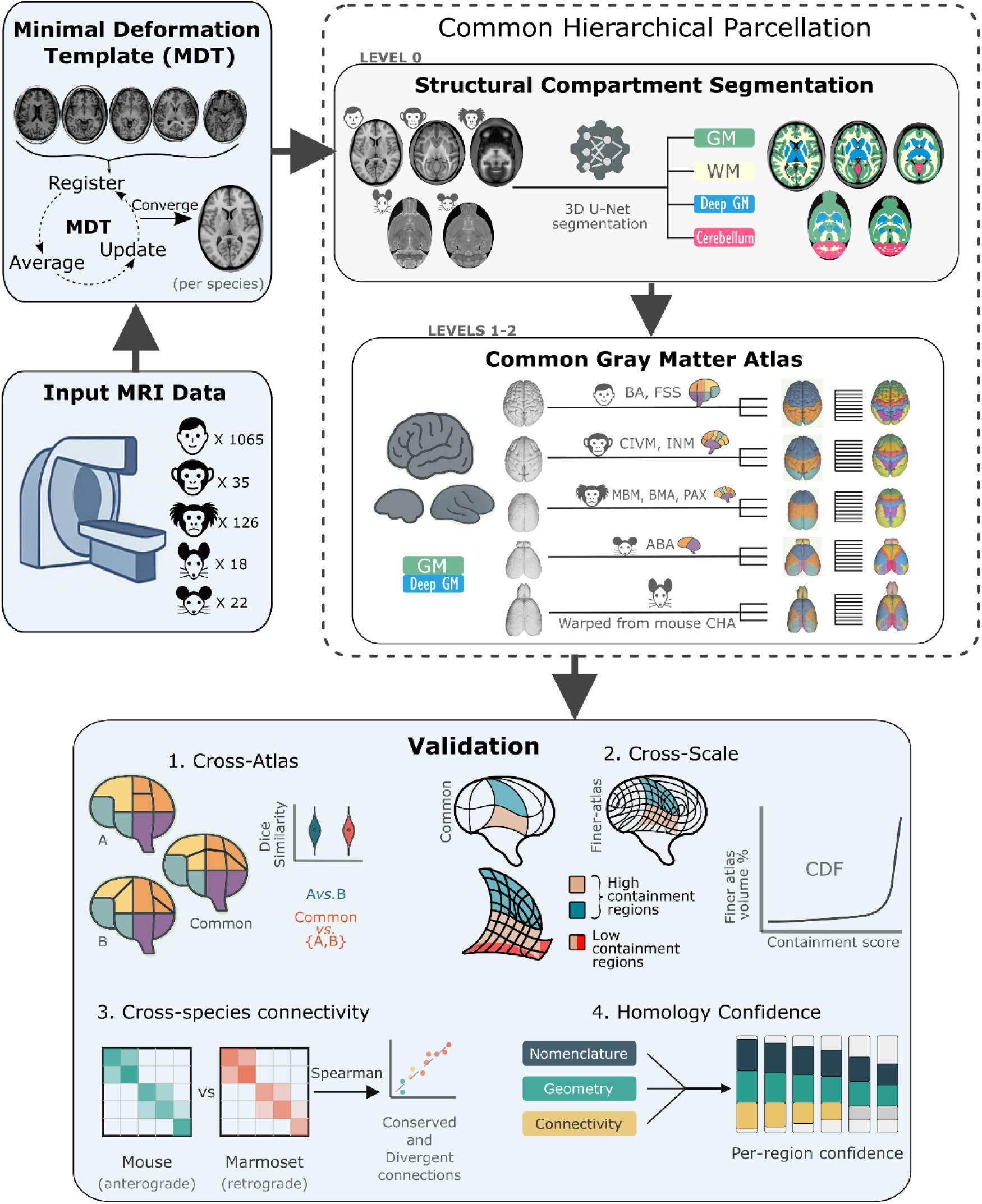
Overview of the common cross-species atlas framework. The workflow integrates species-specific MRI data into a unified anatomical hierarchy through population-based template construction, automated segmentation, and hierarchical parcellation, validated across scales. **(Top Left)** *Input MRI data* were acquired for human, rhesus macaque, marmoset, rat and mouse cohorts. Each species dataset was processed to generate a Minimal Deformation Template (MDT) via iterative diffeomorphic averaging that minimizes anatomical distortion across individuals. **(Top Right)** On each MDT, Level-0 *structural compartment segmentation* was performed using a 3D U-Net to delineate gray matter (GM), white matter (WM), deep gray matter (Deep GM), and cerebellum. These compartment maps provided the foundation for a *Common Hierarchical Parcellation* spanning Levels 1–2. The Common Gray Matter Atlas was then constructed by harmonizing cortical and subcortical parcels across species using literature-guided anatomical homologies (human – FreeSurferSeg/Brodmann; rhesus – CIVM/INM; marmoset – MBM/BMA/Paxinos; mouse – ABA). Rat parcellations were derived by warping the mouse CHA to the rat MDT. **(Bottom)** Validation was performed along four complementary dimensions: (1) Cross-atlas validation quantified overlap (Dice similarity) between CHA and independent human atlases, comparing CHA-to-others with inter-atlas overlaps; (2) Cross-scale containment validation tested how well finer-scale species-specific atlas regions were spatially contained within CHA parcels, with an independent inter-atlas baseline in rat contextualizing boundary correspondence; (3) Cross-species connectivity validation compared independent mouse anterograde and marmoset retrograde tracer datasets mapped to CHA, identifying conserved and divergent connections; (4) A per-region homology confidence index integrated nomenclature directness, cross-species geometric consistency, and tracer connectivity correspondence to quantify the strength of evidence supporting each regional assignment.

Each MDT was then processed using a 3D U-Net segmentation model trained to delineate major tissue classes: gray matter (GM), white matter (WM), deep gray matter (DGM), cerebellum (CBL), and cerebrospinal fluid (CSF). This segmentation defined the Level 0 structural hierarchy, which served as the foundation for all higher-order parcellations. A full list of abbreviations used in this article is provided in Supplementary Table 1.

Based on these tissue-segmented MDTs, we constructed a Common Hierarchical Parcellation to define homologous cortical and subcortical boundaries across species. The framework organizes brain regions into two additional levels: Level 1 macro-regions, which comprise nine major cortical and subcortical divisions, and Level 2 subdivisions, which represent finer-scale anatomical parcels defined by consistent anatomical landmarks and literature-supported homologies. Together, these levels form the Common Gray Matter Atlas, harmonizing parcellations across phylogenetically distant species within a unified coordinate system.

Atlas validation was performed along four complementary dimensions (Fig. 1, bottom). (1) Cross-atlas validation quantified Dice similarity between the common atlas and multiple species-specific atlases, providing a voxelwise measure of regional correspondence relative to existing parcellations. (2) Cross-scale containment validation evaluated the spatial nesting of finer atlas regions within broader common parcels, using containment scores and perturbation analyses to characterize the degree of alignment and boundary correspondence across scales. (3) Cross-species connectivity validation compared independent mouse anterograde and marmoset retrograde tracer datasets mapped to CHA, identifying where connectivity architecture is conserved across species and where it diverges. (4) A per-region homology confidence index integrated nomenclature directness, cross-species geometric consistency, and tracer connectivity correspondence to quantify the strength of evidence supporting each regional assignment.

### 2.2. Cross-Species Parcellation Hierarchy

Figure 2 illustrates the hierarchical parcellation sequence from Level 0 through Level 1 and 2 across the four species. Panels (a–d) depict Level 0 structural compartment segmentation, showing automated 3D U-Net classification of gray matter (GM), white matter (WM), deep gray matter (DGM), and cerebellum (CBL) in human, rhesus, marmoset, and mouse MDTs, respectively. This segmentation establishes the basic tissue architecture on which all higher-order divisions are defined.

**Figure 2.**
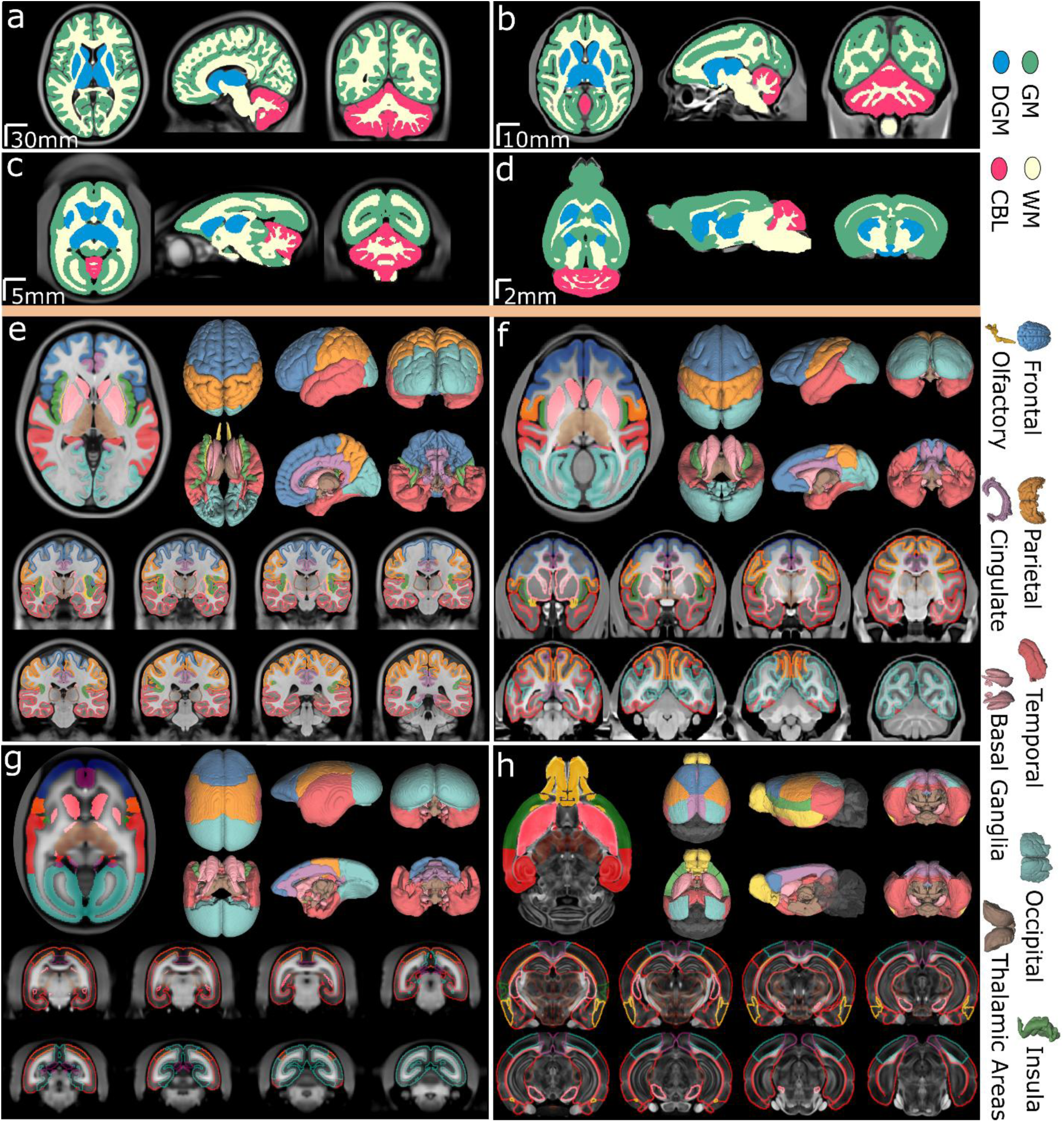
Cross-species construction of the common anatomical atlas. **(a–d)** Tissue segmentation (structural compartments) for human **(a)**, rhesus macaque **(b)**, marmoset **(c)**, and mouse **(d)**. High-resolution T1- and T2-weighted MRI scans were processed using U-NET Studio for automated segmentation into five classes: gray matter (GM, green), white matter (WM, yellow), deep gray matter (DGM, blue), cerebellum (CBL, pink), and cerebrospinal fluid (not shown). Representative sagittal, coronal, and axial views illustrate the structural compartments in each species. **(e–h)** Lobe-level gray matter parcellation (Level-1) for human **(e)**, rhesus **(f)**, marmoset **(g)**, and mouse **(h)**. Nine macro-regions—frontal, parietal, temporal, occipital, insula, cingulate, basal ganglia, thalamus, and olfactory—were defined using consistent anatomical landmarks and human-centric mapping. Cortical regions are displayed with harmonized color-coded boundaries and abbreviations, providing the scaffold for Level-2 subdivisions developed in subsequent analyses. Full mosaic views for all species are given in supplementary figures 1 through 5.

From these structural compartments, the atlas extends to Level 1 macro-regions, representing nine major divisions—frontal (FRO), parietal (PAR), temporal (TEM), occipital (OCC), insular (INS), olfactory (OLF), cingulate (CIN), basal ganglia (BG), and thalamic (THL) systems. These macro-regions, shown across species in Figure 2 (e–h), are defined by consistent anatomical landmarks and human-centric reference mapping while accommodating species-specific anatomical variation.

Figure 3 expands the view to show the full hierarchical organization of Level 1 and 2 structures. GM and DGM compartments form the root of the hierarchy, branching into nine Level 1 divisions and a total of 26 Level 2 subdivisions in primates and 22 in rodents. Harmonized color schemes enable direct cross-species comparison. The parietal (PAR) territory and the caudoputamen remain undivided in rodents (denoted by *), reflecting both true anatomical differences and segmentation limitations. Table 1 provides the full regional definitions, species-specific atlas sources, spatial adjacency criteria, and quantitative geometric consistency scores for all Level 2 regions. This hierarchical framework unifies cortical and subcortical gray matter organization, establishing a scalable anatomical reference for comparative and network-level analyses.

**Figure 3.**
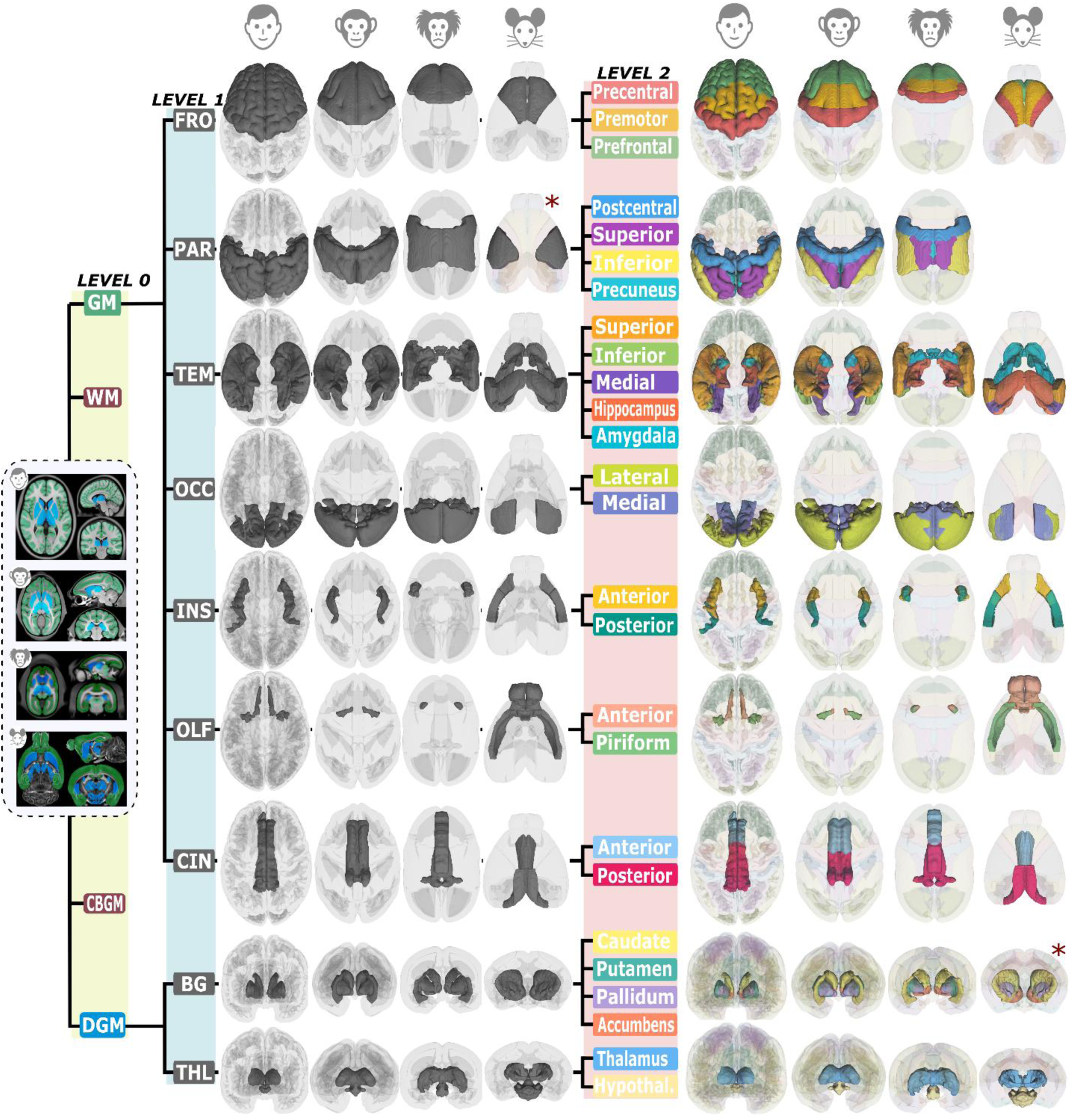
Cross-species parcellation hierarchy. The hierarchical relationships between grey matter parcels are shown across four species (human, rhesus, marmoset, mouse). Four tissue compartments form level-0 of the hierarchy: Grey Matter (GM), White Matter (WM), Cerebellar Grey Matter (CBGM), and Deep Grey Matter (DGM). The GM and DGM compartments form root structures for a total of nine level-1 structures across species (grey). These level-1 structures are further subdivided into a total of 26 level-2 structures in primates and 22 level-2 structures in rodents. Level-1 PAR and level-2 Caudoputamen structures are undivided in rodents (denoted by *). Parcel colors are harmonized across species to facilitate direct visual comparison. Additional sagittal and coronal views of medial structures for all species are given in supplementary figure 6.

**Table 1.** Regional definitions, spatial adjacency criteria, and homology support for each CHA region.

| CHA Region | Definition and coverage | Spatial adjacency<sup>1</sup> | Geo. R<sup>2</sup> | Nom.<sup>3</sup> |
| --- | --- | --- | --- | --- |
| **FRO_Precentral** | Primary motor cortex. Mouse: MOp (ABA). Marmoset: A4ab, A4c (Paxinos). Rhesus: precentral (CIVM). Human: precentral gyrus, BA 4 (FSS/BA). | FRO_Premotor (anterior); PAR (posterior); CIN_Posterior (ventromedial (primates), posteromedial (mouse)) | 0.93 | 2/2/3/3 |
| **FRO_Premotor** | Secondary motor areas. Mouse: MOs (ABA). Marmoset: A6DC, A6DR, A6M, A6Va, A6Vb (Paxinos). Rhesus: premotor (CIVM). Human: premotor cortex, BA 6 (FSS/BA). | FRO_Prefrontal (anteroventral); FRO_Precentral (posterior) | 0.96 | 2/2/3/3 |
| **FRO_Prefrontal** | Mouse: prelimbic, infralimbic, and orbital regions (ABA: PL, ILA, ORB), which serve analogous roles to primate prefrontal cortex but lack laminar and areal complexity (functional analog). Marmoset: A9, A10, A11, A13, A14, A46, A47 (Paxinos). Rhesus: prefrontal (CIVM). Human: dorsolateral, ventrolateral, medial prefrontal, and orbitofrontal areas (FSS/BA). | FRO_Premotor (posterodorsal); CIN_Anterior (medial); OLF_Anterior (posteroventral); OLF_Piriform (posteroventral); INS_Anterior (posteroventrolateral) | 0.98 | 1/2/3/3 |
| **PAR** | Mouse: undivided somatosensory territory, SSp and SSs (ABA). The mouse PAR parcel corresponds primarily to its continuous somatosensory cortex, whereas primates possess a subdivided parietal cortex. PAR is retained as a single undivided territory in rodents to preserve lobe-level comparability. Marmoset: PE, PEC, PF, PFG, PG, PGM, AIP, LIP, MIP, VIP, OPt (Paxinos). Rhesus: parietal (CIVM). Human: postcentral, superior parietal, inferior parietal, precuneus (FSS). Undivided in rodents. | FRO_Precentral (anterior); TEM_Superior (ventrolateral); INS_Posterior (ventrolateral); OCC_Lateral (posteroventral); OCC_Medial (posteromedial); CIN_Posterior (medial) | 0.92 | 2/3/3/3 |
| **TEM_Superior** | Mouse: primary and secondary auditory areas (ABA: AUDp, AUDd, AUDpo, AUDv) (functional analog based on auditory function). Marmoset: Core, BeltL, BeltM, PBC, PBR, STR, FSTR, FSTC, TPOC, TPOR, TPPro, PGB, MST, V5R, V5C, V4L (MBM). Rhesus: superior | PAR (dorsomedial); TEM_Inferior (posteroventral); | 0.91 | 1/2/3/3 |

<sup>1</sup>Spatial adjacency criteria state the neighboring CHA regions and their directions relative to the focal region, as defined during parcellation. Directions were assigned based on conserved anatomical landmarks and literature-guided spatial relationships. Full per-species pairwise spatial adjacencies, computed independently from the atlas label volumes, are provided in Supplementary Table 2.
<sup>2</sup>Geometric consistency (Geo. R) is the mean resultant length of centroid direction vectors across all four species for each region (averaged over all pairwise R values involving that region; range 0–1, where 1 indicates identical relative positioning across species).
<sup>3</sup>Per-species nomenclature directness score (Mo = mouse, Mz = marmoset, Rh = rhesus macaque, Hu = human): 3 = source atlas uses the same or directly equivalent term; 2 = recognized anatomical equivalent; 1 = functional analog supported by literature; 0 = no direct support.

| CHA Region | Definition and coverage | Spatial adjacency<sup>1</sup> | Geo. R<sup>2</sup> | Nom.<sup>3</sup> |
| --- | --- | --- | --- | --- |
|  | temporal (INM). Human: superior and middle temporal gyri (FSS); auditory and auditory association cortex; language/auditory functions emphasized. | INS_Posterior (medial);<br>OCC_Lateral (posterodorsal) |  |  |
| **TEM_Inferior** | Mouse: entorhinal cortex (ABA: ECT), which lacks the laminar complexity of primate IT but shares connectivity properties including input from ventral visual areas and sensitivity to object-level features<sup>13,14</sup> (functional analog). Marmoset: TE1, TE2, TE3, TEO (Paxinos). Rhesus: inferior temporal (INM). Human: inferior temporal gyrus (FSS). | TEM_Superior (anterodorsal);<br>TEM_Medial (posteromedial) | 0.93 | 1/2/3/3 |
| **TEM_Medial** | All species: rhinal/parahippocampal complex. Mouse: entorhinal, perirhinal, postrhinal cortex (ABA: ENTI, ENTm, PERI, POST). Marmoset: Ent, PrhR, PrhC, PHG (Paxinos). Rhesus: medial temporal (INM). Human: entorhinal and parahippocampal cortex (FSS). Supports hippocampal-neocortical interactions. | TEM_Amygdala (anteromedial);<br>TEM_Hippocampus (anterodorsal);<br>TEM_Inferior (anterolateral);<br>OCC_Lateral (posterodorsal) | 0.95 | 3/3/3/3 |
| **TEM_Hippocampus** | Conserved structure across species. Mouse: HPF (ABA). Marmoset: HipF (Paxinos/MBM). Rhesus: hippocampus (CIVM). Human: hippocampus (FSS). | TEM_Amygdala (anteroventral);<br>TEM_Medial (posteroventral);<br>CIN_Posterior (dorsomedial);<br>OCC_Medial (posterodorsal);<br>THL_Thalamus (dorsomedial) | 0.92 | 3/3/3/3 |
| **TEM_Amygdala** | Preserved across species; limbic/olfactory interfaces. Mouse: AMY (ABA). Marmoset: Amy (Paxinos/MBM). Rhesus: amygdala (CIVM). Human: amygdala (FSS). | TEM_Hippocampus (posterodorsal);<br>TEM_Medial (posterolateral);<br>BG_CaudoPutamen (anterodorsal) | 0.97 | 3/3/3/3 |
| **OCC_Lateral** | Mouse: lateral visual areas (ABA: VISl, VISal, VISrl, VISa, VISli). Marmoset: V2L, V3L, V4, V4T, V5 (Paxinos). Rhesus: lateral occipital (INM). Human: superior, middle, inferior occipital gyri (FSS). | PAR (anterodorsal);<br>TEM_Superior (anteroventral);<br>TEM_Medial (anteroventral);<br>OCC_Medial (anteromedial) | 0.96 | 2/2/3/3 |
| **OCC_Medial** | Mouse: primary and medial visual areas (ABA: VISp, VISm, VISpm). Marmoset: V1, V2M, A19M (Paxinos). Rhesus: medial occipital (INM). Human: lingual gyrus, cuneus, calcarine (FSS). | CIN_Posterior (anterior);<br>PAR (anterodorsal);<br>TEM_Hippocampus (anteroventral);<br>OCC_Lateral (posterolateral) | 0.94 | 2/2/3/3 |
| **INS_Anterior** | Agranular insula with adjacency to piriform emphasized. Mouse: agranular insular cortex (ABA: AI). Marmoset: AI (Paxinos). Rhesus: anterior insula (CIVM). Human: anterior insula (FSS). | FRO_Prefrontal (anterodorsal);<br>INS_Posterior (posterior);<br>OLF_Piriform (ventromedial) | 0.92 | 2/2/3/3 |
| **INS_Posterior** | Granular insula; consistent across primates. Mouse: gustatory and visceral areas (ABA: GU, VISC). Marmoset: GI, DI (Paxinos). Rhesus: posterior insula (CIVM). Human: posterior insula (FSS). | INS_Anterior (anterior); PAR (posterodorsal); TEM_Superior (lateral) | 0.88 | 2/2/3/3 |
| **OLF_Anterior** | Anteromedial adjacency to piriform in all species; routes olfactory inputs to piriform. Olfactory bulb included in rodents and human. Mouse: MOB, AOB, AON (ABA). Marmoset: limited coverage (Paxinos). Rhesus: limited coverage (CIVM). Human: olfactory (FSS). Landmark-guided in primates. | FRO_Prefrontal (anterodorsal); OLF_Piriform (posteroventral) | 0.92 | 3/2/2/3 |
| **OLF_Piriform** | Piriform cortex preserved as separate region; adjacency to insula. Mouse: PIR (ABA). Marmoset: Pir (Paxinos). Rhesus: piriform (CIVM). Human: piriform (FSS). | OLF_Anterior (anteromedial); FRO_Prefrontal (anterodorsal); INS_Anterior (dorsolateral) | 0.95 | 3/3/3/3 |
| **CIN_Anterior** | Anterior cingulate. Anterior vs posterior split supported by cytoarchitecture and connectivity<sup>15</sup>. Mouse: anterior cingulate area (ABA: ACA). Marmoset: A24a-d, A32, A32V, A25 (Paxinos). Rhesus: anterior cingulate (CIVM). Human: anterior cingulate, BA 24/32 (FSS/BA). | CIN_Posterior (posterodorsal); FRO_Prefrontal (lateral) | 0.92 | 2/2/3/3 |
| **CIN_Posterior** | Posterior cingulate/retrosplenial. Mouse: retrosplenial cortex (ABA: RSPagl, RSPd, RSPv). In mouse, retrosplenial cortex occupies the dorsal medial surface adjacent to PAR, whereas in primates it is located on the medial wall. Marmoset: A23a-c, A29a-c, A29d, A30, A31 (Paxinos). Rhesus: posterior cingulate (CIVM). Human: posterior cingulate, BA 23/29/30/31 (FSS/BA). | CIN_Anterior (anteroventral); FRO_Precentral (anterolateral); PAR (lateral); OCC_Medial (posterior); TEM_Hippocampus (ventrolateral) | 0.96 | 2/2/3/3 |
| **BG_CaudoPutamen** | Core striatal component. Subdivided into caudate + putamen in primates; undivided in rodents due to absence of clear anatomical boundary in mouse MRI (Allen Mouse Brain Atlas likewise treats caudoputamen as a single complex). Mouse: CP (ABA). Marmoset: Cd, Pu (Paxinos/MBM). Rhesus: caudate, putamen (CIVM). Human: caudate, putamen (FSS). | BG_Accumbens (anteroventral); BG_Pallidum (posteroventral); TEM_Amygdala (posteroventral) | 0.94 | 3/3/3/3 |
| **BG_Accumbens** | Kept distinct for translational relevance. Mouse: ACB (ABA). Marmoset: Acb (Paxinos/MBM). Rhesus: accumbens (CIVM). Human: accumbens (FSS). | BG_CaudoPutamen (posterodorsal) | 0.95 | 3/3/3/3 |
| **BG_Pallidum** | Globus pallidus; robust segmentation across species. Mouse: PAL (ABA). Marmoset: GP (Paxinos/MBM). Rhesus: pallidum (CIVM). Human: pallidum (FSS). | BG_CaudoPutamen (anterodorsal) | 0.95 | 3/3/3/3 |
| **THL_Thalamus** | Well-defined across species; treated separately from basal ganglia. Mouse: TH (ABA). Marmoset: Thal (Paxinos/MBM). Rhesus: thalamus (CIVM). Human: thalamus (FSS). | TEM_Hippocampus (ventrolateral) | 0.92 | 3/3/3/3 |
| **THL_Hypothalamus** | Well-defined across species. Mouse: HY (ABA). Marmoset: Hypo (Paxinos/MBM). Rhesus: hypothalamus (CIVM). Human: hypothalamus (FSS). | THL_Thalamus (dorsal);<br>TEM_Amygdala (ventrolateral) | 0.95 | 3/3/3/3 |
| **Excluded** | Clastrum, substantia nigra, subthalamic nucleus: homologous in principle but excluded due to MRI segmentation limitations across species. | — | — | — |

**Abbreviations:** ABA = Allen Brain Atlas (mouse); Paxinos = Paxinos stereotaxic atlas (marmoset); CIVM = Center for In Vivo Microscopy atlas (rhesus); INM = Inia19-NeuroMaps (rhesus); MBM = Marmoset Brain Mapping atlas; FSS = FreeSurferSeg (human); BA = Brodmann areas (human). Nomenclature scores: Mo = mouse; Mz = marmoset; Rh = rhesus macaque; Hu = human.

### 2.3. Volumetric Composition Across Species

Figure 4 quantifies species differences in volumetric organization across Level 2 regions. Panel (a) presents dumbbell plots of normalized Level 2 volume fractions within each species. Volumetric fractions revealed clear differences across species MDTs in cortical and subcortical regions (Fig. 4a, b). The human MDT showed the largest proportional volumes in prefrontal (0.22) and parietal (0.19) cortex, consistent with well-established expansion of association cortices in primates and especially in humans^16–18^. The rhesus macaque MDT also showed elevated prefrontal (0.15) and superior temporal (0.16) fractions, in agreement with comparative MRI and anatomical studies highlighting enlarged association and temporal systems in Old World primates^19,20^.

**Figure 4.**
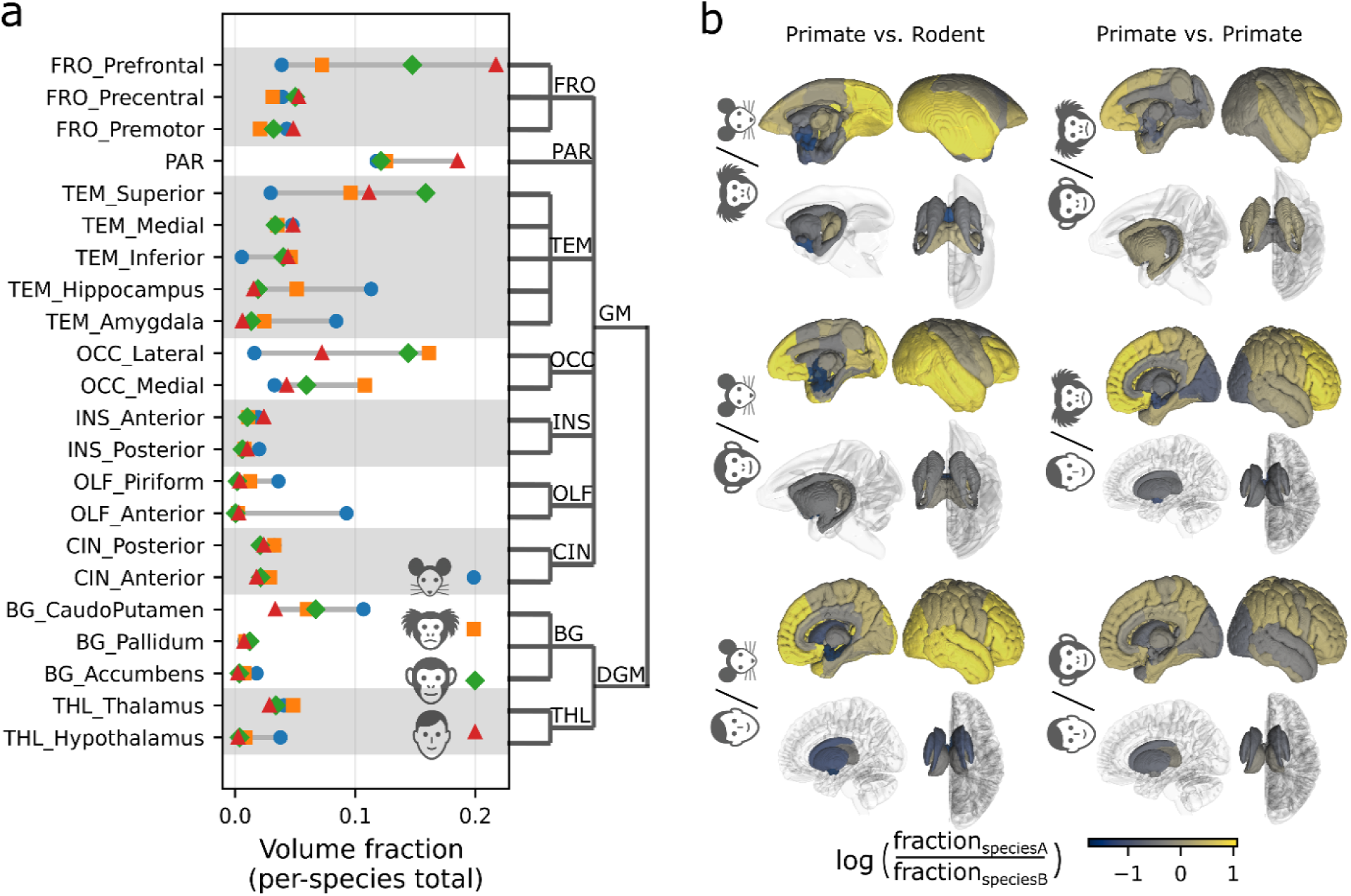
Volumetric composition across species MDTs. **(a)** Dumbbell plot of Level-2 volume fractions (the proportion of each region’s volume relative to the total parcellated gray matter volume within each species MDT). For each region, colored markers denote species values. **(b)** Pairwise 3D maps of relative volumetric investment for six interspecies comparisons. Colors encode log(*fraction*_*species A*_⁄*fraction*_*species B*_); positive values indicate greater fractional investment in species A. For each comparison, color is mapped onto the species A brain template. Because these measurements are derived from population-averaged minimal deformation templates rather than individual subjects, they reflect species-level descriptive properties and do not capture individual variability.

The marmoset MDT exhibited comparatively higher fractions in lateral and medial occipital cortex (0.16 and 0.11), as well as appreciable superior temporal volume (0.10), consistent with their relatively extensive visual and multisensory cortical territories^21,22^. In contrast, the mouse MDT showed the highest proportional volumes in hippocampus (0.11), amygdala (0.08), anterior olfactory regions (0.09), and piriform cortex (0.036), matching the prominence of allocortical and olfactory systems described in rodents^23,24^. Across species, deep gray-matter regions, including pallidum, accumbens, and thalamus, varied more modestly in fractional volume than the cortical systems, in line with general principles of coordinated but constrained scaling across brain structures described in comparative mammalian neuroanatomy^25,26^. Panel (b) displays pairwise 3D maps of relative volumetric investment, color-coded by the logarithmic ratio of fractional volumes between species. These maps reveal cortical expansion in the human MDT and stronger limbic weighting in the rodent MDT, reflecting both conserved and lineage-specific scaling principles. Together, these results indicate that the common atlas enables comparison of homologous regional definitions and highlights volumetric allocation differences across MDTs that are consistent with primate associative enlargement and rodent limbic specialization.

### 2.4. Validation of the Common Atlas

We evaluated the anatomical fidelity of the common atlas through four complementary approaches: (i) cross-atlas Dice similarity against established human parcellations and mouse ABA major divisions, (ii) containment validation against finer-scale species-specific atlases, (iii) cross-species connectivity validation using independent mouse and marmoset invasive tracer datasets, and (iv) a per-region homology confidence index integrating nomenclature directness, geometric consistency, and connectivity correspondence.

#### 2.4.1. Validation of the Common Human Atlas Using Cross-Atlas Dice Similarity

The common human atlas showed broadly consistent alignment with standardized Neuroparc^4^ parcellations. Most Level-2 regions matched 4–8 labels across reference atlases (e.g., hippocampus: 8; precuneus: 6; nucleus accumbens: 2). Regions such as the hypothalamus and finer olfactory subdivisions were not consistently labeled or directly mappable. However, all Level-2 parcels were evaluated where correspondence could be established.

Violin plots of Dice similarity demonstrated that Common vs. Others typically achieved higher overlap than Others vs. Others, indicating greater internal consistency of the common atlas relative to inter-atlas variability (Fig. 5a). Scatterplots of ΔMedian Dice versus absolute Dice confirmed that nearly all regions fell above zero (median Δ = +0.125, Wilcoxon signed-rank one-sided p = 1.22 × 10⁻⁴), indicating that CHA does not introduce additional inconsistency beyond what already exists among established human atlases. Cortical regions displayed greater variability than subcortical structures such as the thalamus and putamen, consistent with prior benchmarks reporting poor cortical but stronger subcortical concordance across human atlases^1–4^. Given these well-documented inconsistencies, no single parcellation can serve as ground truth for cortical boundary definitions, and that CHA median Dice consistently met or exceeded inter-atlas agreement is noted as an encouraging but secondary observation.

**Figure 5.**
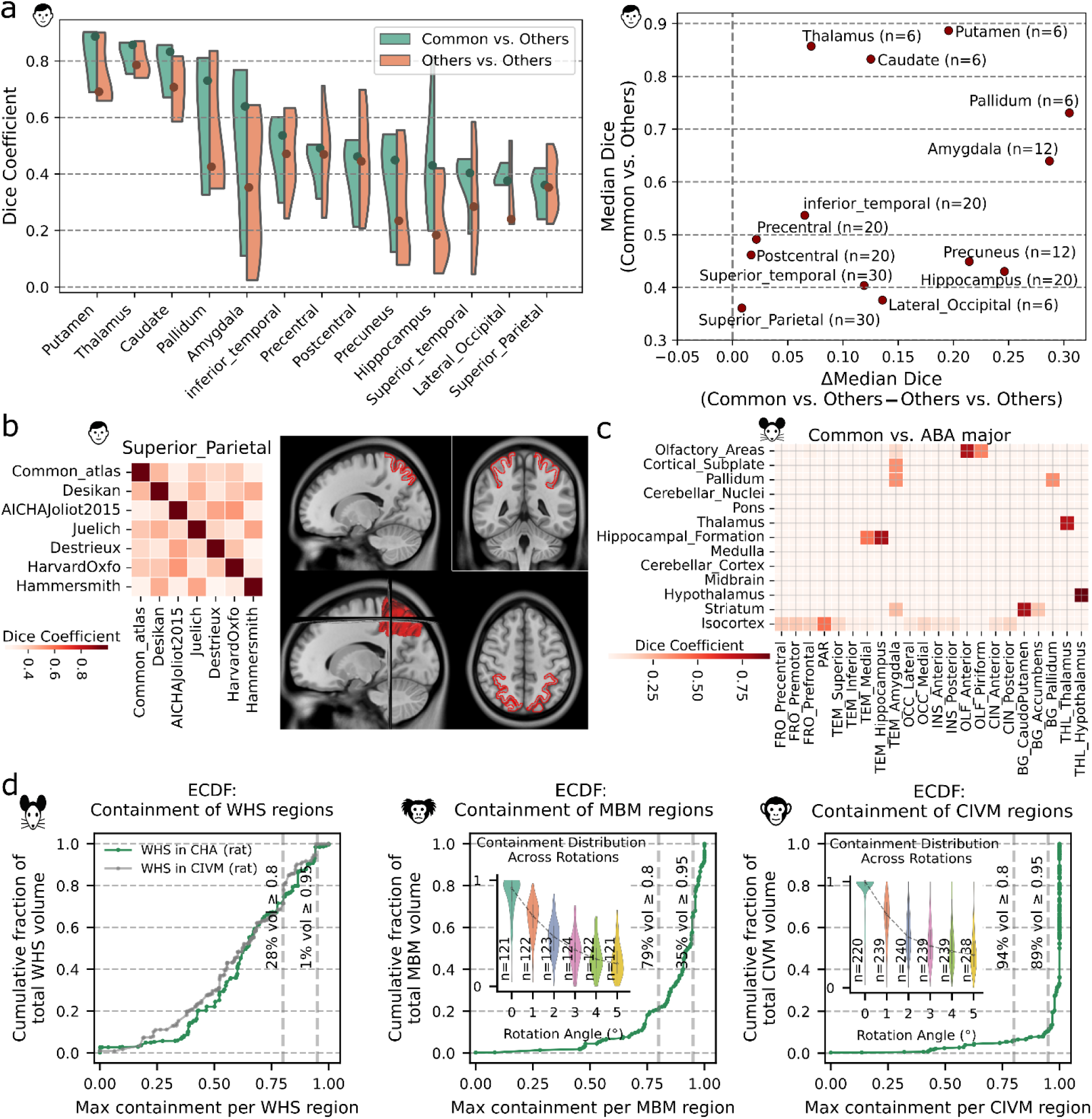
Cross-Atlas validation of the common framework. **(a)** Validation of human parcellation using cross-atlas Dice similarity. Dice coefficients were computed for each common atlas region against Neuroparc references^4^ and compared to overlaps among non-common atlases ("Common vs. Others" refers to overlap between CHA and each Neuroparc atlas; "Others vs. Others" refers to overlap among Neuroparc atlases themselves, excluding CHA). Both CHA and the Neuroparc atlases are defined in ICBM152 standard space, requiring no inter-atlas registration; the CHA human atlas (0.5 mm isotropic) was downsampled to match Neuroparc resolution (1 mm isotropic). Violin plots show region-wise distributions; scatterplots summarize median Dice and ΔMedian differences. CHA achieved higher median Dice than inter-atlas agreement in all 13 regions evaluated (median Δ = +0.125, Wilcoxon signed-rank one-sided p = 1.22 × 10⁻⁴), confirming that the common atlas does not introduce additional boundary inconsistency relative to existing inter-atlas variability. **(b)** Expanded analysis of superior parietal lobule, the weakest Dice performer in panel a. Left: Dice matrix comparing the common SPL to six Neuroparc atlases. Right: common SPL overlaid on T1w MRI, showing alignment with gyral boundaries despite low inter-atlas concordance. **(c)** Dice similarity between the 13 major divisions of the Allen Brain Atlas (ABA) and common Level-2 regions. High Dice values occurred where there was direct one-to-one correspondence between ABA and CHA (e.g., thalamus, hypothalamus, olfactory, hippocampus). Lower Dice values for broad ABA parcels such as isocortex reflect the intentional subdivision of mouse cortical territory into human-centric divisions to enable cross-species comparison. **(d)** Cross-species containment validation. Volume-weighted empirical cumulative distribution functions (ECDFs) of maximum containment scores for finer-scale species-specific atlas regions within CHA parcels. Left: Rat Waxholm Space atlas (WHS)^27^ evaluated against both CHA (green) and the Duke CIVM Wistar atlas^28^ (gray) as an independent inter-atlas baseline, showing that CHA containment is comparable to agreement between two independent rat atlases. Center: Marmoset Brain Mapping atlas (MBM) regions in CHA, with 79% of volume achieving containment ≥ 0.80 and 35% ≥ 0.95. Right: CIVM rhesus macaque atlas regions in CHA, with 94% of volume achieving containment ≥ 0.80 and 89% ≥ 0.95. Insets for marmoset and rhesus show systematic decay of containment scores under rotational perturbation, confirming sensitivity of the metric to spatial disruption. Mouse containment (perfect by construction, as ABA served as the sole source for CHA mouse labels) is shown in Supplementary Figure 7. The gradient from rat to marmoset to rhesus reflects differences in the available species-specific atlas infrastructure rather than CHA boundary quality.

Certain cortical regions, such as the superior parietal lobule (SPL), showed lower correspondence (median Dice ≈ 0.37), although the common atlas still performed slightly better than inter-atlas comparisons. The expanded Dice matrix (Fig. 5b) confirmed low similarity across all atlases, highlighting the variability of SPL boundaries in existing human parcellations. However, overlaying the common SPL on T1w MRI demonstrated clear alignment with the superior parietal gyrus, supporting the anatomical validity of the common definition despite poor cross-atlas agreement.

#### 2.4.2. Evaluation Against Mouse ABA Major Regions

Dice similarity analyses revealed two complementary patterns. For ABA regions with direct correspondence to common parcels, such as hypothalamus, thalamus, olfactory areas, and hippocampus, Dice values were high, reflecting strong spatial alignment (Fig. 5c). In contrast, broad ABA categories like *Isocortex* showed lower values because these regions were intentionally subdivided across multiple common ROIs to match human-centric definitions. This outcome indicates that the common framework preserves homologous subcortical structures with high fidelity, while deliberately enforcing finer cortical subdivisions to support cross-species consistency.

#### 2.4.3. Containment Validation Against Finer-Scale Species-Specific Atlases

Containment validation was used to benchmark anatomical correspondence between finer-scale species-specific atlases and the common framework (Fig. 5d). The containment metric measures the fraction of each finer ROI that is fully contained within its corresponding common parcel, thereby quantifying how well the broader common regions align with and respect the boundaries of more detailed species-specific parcellations. This provides a direct test of whether the common atlas captures homologous anatomical divisions without overstepping finer boundaries.

Across species, validation confirmed strong correspondence between common parcels and species-specific atlases. In rhesus, compared with the CIVM atlas (220 regions), 94% of volume achieved a containment score ≥ 0.80 and 89% ≥ 0.95, demonstrating high fidelity to established boundaries. In marmoset, compared with the MBM atlas (122 regions), 79% of volume achieved ≥ 0.80 and 35% ≥ 0.95. In rat, compared with the Waxholm Space atlas^27^ (133 regions), 28% of volume achieved ≥ 0.80 and 1% ≥ 0.95. The WHS atlas is based on a single Sprague Dawley specimen, whereas our rat MDT was constructed from 18 Wistar rats, introducing a strain difference that may contribute to boundary misalignment. To contextualize these values, we computed an independent inter-atlas baseline by evaluating containment of WHS regions within the Duke Wistar atlas (54 regions)^28^, yielding 31% of volume ≥ 0.80 and 0% ≥ 0.95. CHA containment was thus comparable to agreement between two independent rat atlases, indicating that boundary correspondence is limited by the available rat atlas infrastructure rather than by CHA itself. The rat atlas landscape and its implications for CHA validation are discussed further in Section 3.6.

To test the sensitivity of the containment metric, common labels were rigidly rotated in fixed angular increments and containment scores recomputed for each configuration (Fig. 5d insets, marmoset and rhesus; Supplementary Figure 7, mouse). Containment declined systematically with increasing rotation, confirming the metric’s sensitivity to spatial disruption. Together with the perturbation analysis, these results demonstrate that containment scores reflect genuine anatomical alignment rather than trivial overlap.

The minority of regions with low containment reflect deliberate design choices. Where a primary atlas (e.g., MBM) did not provide clear homology, secondary resources (e.g., Paxinos, Brain/MINDS) were used to reach cross-species consistency. Given the well-documented inconsistencies among published atlases, occasional low containments are expected and reflect our emphasis on preserving biologically plausible, literature-supported homologies rather than enforcing strict one-to-one correspondence with any single atlas.

#### 2.4.4. Cross-Species Connectivity validation

To validate the CHA’s regional definitions using independent connectivity data, we compared mouse and marmoset corticocortical connectivity matrices mapped to CHA (Fig. 6). Mouse connectivity was derived from ∼1,197 anterograde viral tracing experiments (Allen Mouse Brain Connectivity Atlas)^29^, while marmoset connectivity was derived from 143 retrograde fluorescent tracer injections (Marmoset Brain Connectivity Atlas)^30^, mapped to CHA via Paxinos-to-CHA containment fractions. Despite differences in tracing methodology (anterograde vs. retrograde), species, and over an order-of-magnitude difference in dataset scale, the two matrices showed significant rank-order correspondence across 101 directed corticocortical connections (Spearman ρ = 0.61, p < 10⁻¹¹; Fig. 6c). Agreement increased with marmoset tracer coverage (quartile mean agreement: Q1 = 0.76, Q4 = 0.84; Fig. 6c, inset), indicating that correspondence strengthens where both datasets are well sampled. Sorted per-connection agreement values are provided in Supplementary Table 3. A more extensive connectivity-based validation across all four species using diffusion MRI is provided in^31^.

**Figure 6.**
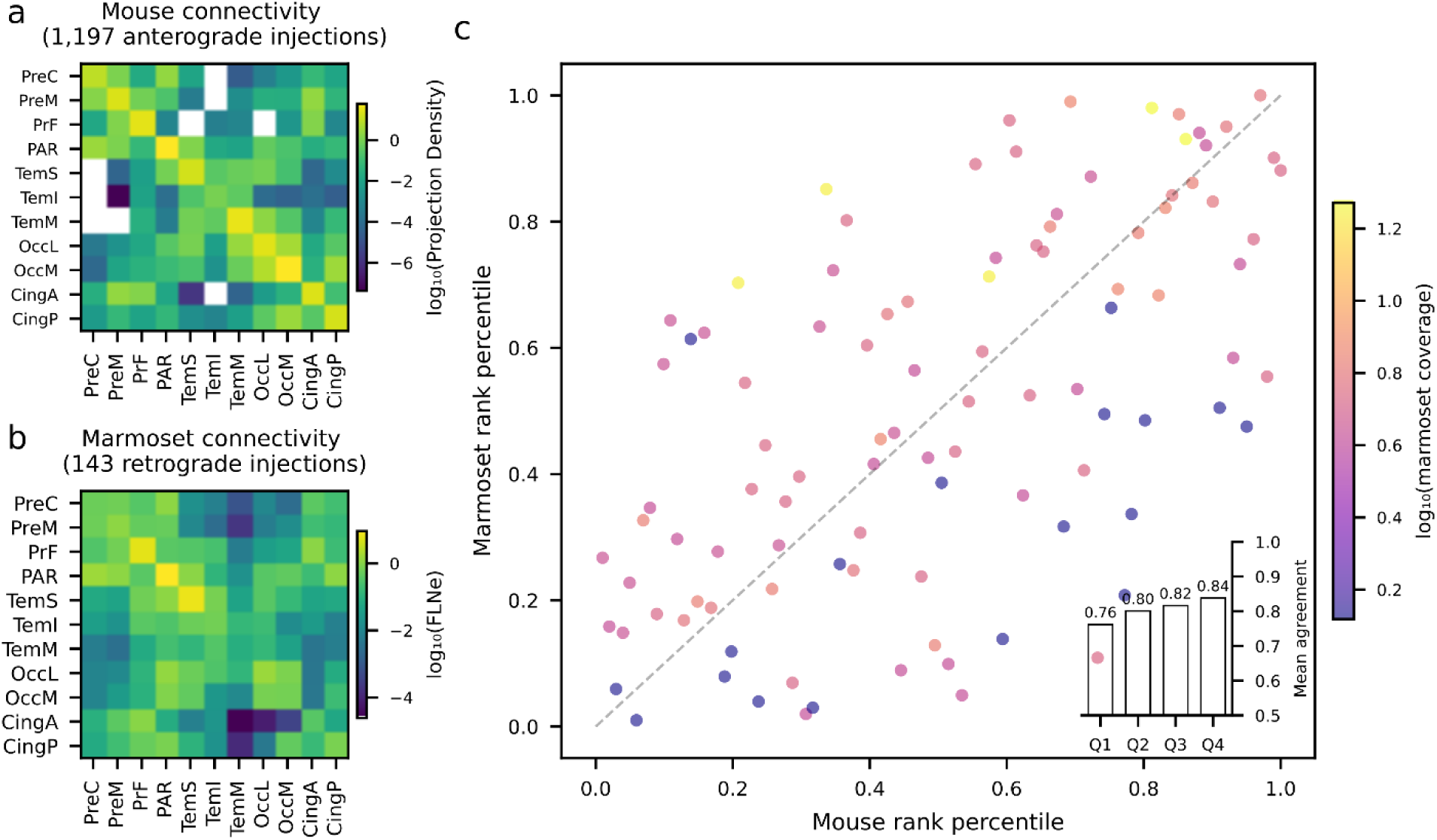
Cross-species connectivity validation using independent invasive tracer datasets. **(a)** Mouse corticocortical connectivity matrix derived from ∼1,197 anterograde viral tracing experiments (Allen Mouse Brain Connectivity Atlas)^29^, mapped to the common hierarchical atlas (CHA). Values represent directed projection densities (log₁₀-scaled) computed from injection and projection coverage fractions summed across experiments. **(b)** Marmoset corticocortical connectivity matrix derived from 143 retrograde fluorescent tracer injections (Marmoset Brain Connectivity Atlas)^30^, mapped to CHA via Paxinos-to-CHA containment fractions. Values represent the fraction of extrinsic labeled neurons (FLNe, log₁₀-scaled). Four cortical CHA regions lacking systematic marmoset injection coverage (INS_Anterior, INS_Posterior, OLF_Anterior, OLF_Piriform) were excluded from the analysis. **(c)** Rank-percentile comparison of mouse and marmoset directed connection weights for all region pairs where both matrices have nonzero entries (n = 101; Spearman ρ = 0.61, p < 10⁻¹¹). Each point represents a directed connection between two CHA regions; color indicates marmoset tracer coverage (log₁₀ of the minimum total signal across the two regions in the pair). Inset: mean rank-percentile agreement (1 minus the absolute difference in rank percentiles) grouped by marmoset coverage quartile, showing increasing correspondence with tracer coverage (Q1 = 0.76, Q4 = 0.84).

These tracer-based results are consistent with independent cross-species comparisons reporting greater conservation in sensorimotor systems and greater divergence in association cortex, including transcriptomic analyses^32^, functional connectivity^33^, white matter connectivity blueprints^34,35^, and directed structural connectomics^31^. This convergence across modalities provides further support for the biological validity of CHA’s regional definitions.

Together, these comparisons reveal a systematic pattern of cross-species organization that extends beyond validation of the atlas itself. Connections linking primary sensorimotor regions (e.g., precentral–parietal) exhibit consistently high correspondence between mouse and marmoset, indicating strong conservation of sensorimotor circuitry across species. In contrast, connections involving association territories, particularly prefrontal and inferior temporal regions, show substantially lower correspondence, reflecting divergence in higher-order integrative systems. This conserved–divergent gradient is not imposed by the parcellation but emerges from independent tracer datasets mapped into a common coordinate system. As such, the common atlas enables a quantitative delineation of where cross-species correspondence is strong versus where it breaks down, providing a principled basis for interpreting the translational relevance of circuit-level findings across model systems.

#### 2.4.5. Per-region Homology Confidence

To quantify the strength of evidence supporting each CHA regional assignment, we developed a composite homology confidence index integrating three independent criteria (Figure 7).

**Figure 7.**
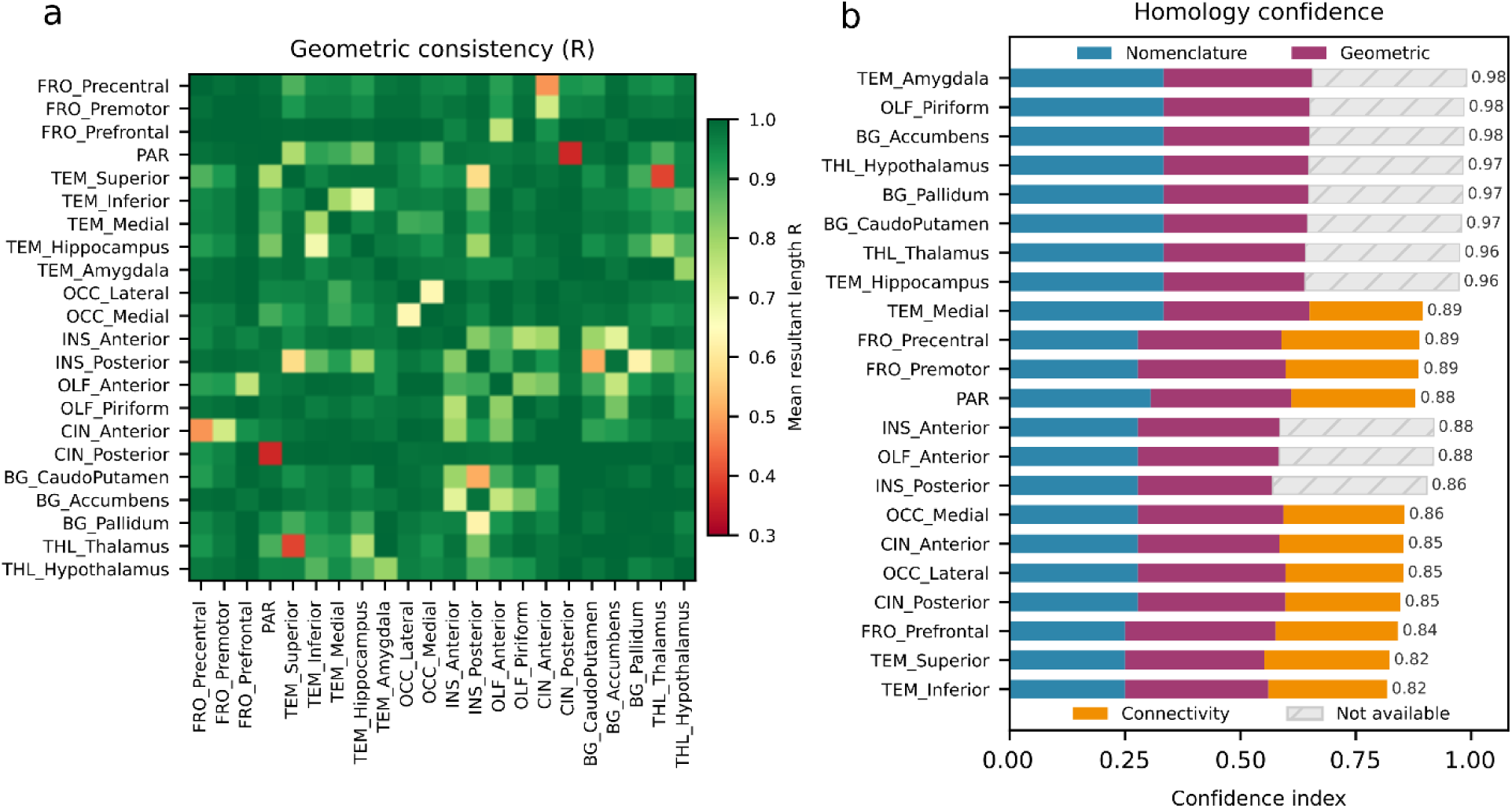
Pairwise geometric consistency and per-region homology confidence index. **(a)** Pairwise geometric consistency (mean resultant length R) of centroid direction vectors across all four species for each CHA region pair. R = 1 indicates identical relative spatial positioning across species; lower values indicate cross-species displacement of the spatial relationship. The lowest consistency is observed for PAR-CIN_Posterior (R = 0.36), reflecting the known shift of retrosplenial cortex from the dorsal surface in rodents to the medial wall in primates. **(b)** Composite homology confidence index integrating three independent criteria, each contributing equally (up to one-third): nomenclature directness (blue; 0-3 scale per species based on atlas support for the CHA assignment, averaged and normalized), geometric consistency (magenta; per-region mean of pairwise R values from panel a), and cross-species connectivity correspondence (orange; per-region mean rank-percentile agreement between mouse anterograde and marmoset retrograde tracer connectivity mapped to CHA). Combined index shown at right of each bar. Hatched gray segments indicate criteria unavailable for evaluation: connectivity could not be assessed for four cortical regions lacking marmoset injection coverage (INS_Anterior, INS_Posterior, OLF_Anterior, OLF_Piriform) or for subcortical regions outside the corticocortical tracer dataset. Regions sorted by combined confidence (ascending from bottom).

Nomenclature directness (Table – 1) was highest (normalized score = 1.0) for regions with explicit, consistent labels across all species-specific source atlases, including hippocampus, amygdala, piriform cortex, medial temporal cortex, and all subcortical regions. The lowest scores (0.75) were assigned to FRO_Prefrontal, TEM_Superior, and TEM_Inferior, reflecting reliance on functional analogy for the mouse assignments, most notably the mapping of mouse ectorhinal cortex to TEM_Inferior.

Geometric consistency, quantified by the mean resultant length (R) of centroid direction vectors across species^36^, was high for all CHA regions (range: 0.878-0.979; Fig. 7a). The highest consistency was observed for FRO_Prefrontal (R = 0.979), indicating a highly conserved spatial position relative to all other regions despite substantial differences in prefrontal complexity across species. The lowest was observed for INS_Posterior (R = 0.878), driven by species differences in the spatial relationship between posterior insula and temporal cortex, caudoputamen and pallidum (Fig. 7a): in rodents, temporal cortex lies purely posterior to the insula, whereas in primates it extends anteriorly to wrap around the entire insular region, substantially altering the relative geometry. At the pair level, the lowest geometric consistency was between CIN_Posterior and PAR (R = 0.360), reflecting the known shift of retrosplenial cortex from the dorsal surface in rodents, where it is directly adjacent to parietal somatosensory cortex, to the medial wall in primates due to cortical folding. TEM_Superior and THL_Thalamus (R = 0.395) also showed low pairwise consistency, reflecting species differences in how temporal cortex wraps around subcortical structures. A complete set of spatial adjacencies are provided in Supplementary Table 2 for each species.

Cross-species connectivity correspondence, assessed through the mouse-marmoset tracer comparison (Fig. 6), was available for 11 cortical CHA regions with adequate marmoset injection coverage. Per-region mean agreement ranged from 0.733 (TEM_Medial) to 0.899 (FRO_Precentral), with sensorimotor regions showing the strongest correspondence and temporal association regions the weakest, consistent with the broader pattern of sensorimotor conservation and association divergence observed across modalities^32,33^.

The combined homology confidence index ranged from 0.818 (TEM_Inferior) to 0.985 (TEM_Amygdala; Fig. 7b). Regions with the highest combined confidence (>0.95) were those with explicit nomenclature across all species and either strong connectivity correspondence or consistent geometric relationships: amygdala, piriform, accumbens, hypothalamus, pallidum, caudoputamen, thalamus, and hippocampus. The lowest confidence (0.82-0.85) was associated with cortical regions where mouse assignments required functional analogy (TEM_Superior, TEM_Inferior) or where cross-species connectivity divergence was greatest (CIN_Posterior, FRO_Prefrontal). Four cortical regions (INS_Anterior, INS_Posterior, OLF_Anterior, OLF_Piriform) and all subcortical regions could not be evaluated on the connectivity criterion due to the cortical scope and incomplete coverage of the marmoset tracer dataset; these are marked in Fig. 7b and represent targets where future primate tracing experiments would most improve confidence.

## 3. Discussion

### 3.1. A consistent hierarchical parcellation and cross-species insights

In this study, we introduce a standardized anatomical framework that aligns cortical and subcortical parcellations across rodents, nonhuman primates, and humans. Population-averaged minimal deformation templates (MDTs) were constructed for each species to provide anatomically consistent reference spaces for cross-species alignment. While humans and marmosets already had established MDTs (ICBM2009a and Brain/MINDS, respectively), we generated new MDTs for mouse and rhesus macaque using high-resolution MRI and iterative diffeomorphic averaging. To ensure consistency across species, we applied a unified 3D U-Net segmentation network to classify tissue compartments into gray matter, white matter, deep nuclei, cerebellum, and CSF, providing a robust and reproducible baseline for higher-level parcellations. This strategy reduces variability introduced by manual delineation or heterogeneous processing methods and ensures that all subsequent parcellations are grounded in a consistent tissue definition. Building on these MDTs and tissue segmentations, we delineated a common hierarchical atlas (CHA) spanning Levels 0 through 2, enabling direct comparison of cortical and subcortical organization across species within a unified coordinate system.

Beyond providing a unified anatomical framework, the present work establishes a quantitative basis for evaluating cross-species correspondence in brain organization. By integrating independent tracer datasets within a common parcellation, we show that conservation across species follows a structured gradient, with sensorimotor systems exhibiting high correspondence and association cortices showing progressive divergence. This pattern is consistent with prior observations from transcriptomic^32^, functional^33^, and connectivity^34,35^ studies, but is here captured within a single anatomically grounded framework that enables direct region-to-region comparison. Importantly, this enables a shift from qualitative assumptions of homology to quantitative assessment: regions and connections can be explicitly evaluated for their degree of cross-species correspondence. In this sense, the common atlas functions not only as a reference space, but as a tool for predicting where findings in model organisms are likely to generalize to primates and humans, and where species-specific specialization should be expected.

### 3.2. Volumetric composition across species MDTs

Our analyses also revealed systematic differences in the volumetric ratios of major brain regions across species MDTs, providing quantitative insights into conserved versus lineage-specific organizational features. The human MDT showed disproportionately larger allocations to frontal and parietal cortices, consistent with expanded associative functions. The rhesus MDT exhibited a higher volume fraction for superior temporal cortex (0.16) than the human MDT (0.08), marmoset MDT (0.10), or mouse MDT (0.06); this reflects the proportional composition of each species’ gray matter rather than absolute temporal size, as the human frontal expansion compresses the relative contribution of other regions. The mouse MDT devoted proportionally more volume to limbic and sensorimotor structures such as hippocampus, amygdala, olfactory regions, and caudoputamen. Despite these divergences, several subcortical and paralimbic structures, including the thalamus, pallidum, accumbens, and insula, demonstrated relatively conserved volumetric scaling across species MDTs. Multivariate approaches examining relative volumes within hierarchical levels, or combinations of volumes to investigate brain-wide compositional trajectories, represent a promising extension of these descriptive analyses.

### 3.3. Validation of cross-species regional definitions

Cross-atlas Dice validation confirmed that CHA achieves spatial overlap with established human parcellations at least equivalent to the agreement those atlases achieve with one another (median Δ = +0.125, Wilcoxon signed-rank p = 1.22 × 10⁻⁴), indicating that CHA does not introduce additional boundary inconsistency beyond what already exists among independently constructed human atlases. For other species, comparably rich sets of overlapping parcellations do not exist, precluding analogous inter-atlas benchmarking. We therefore performed containment validation, quantifying how well finer-scale species-specific atlas regions are spatially contained within CHA parcels. Rhesus containment was highest (94% of volume at score ≥ 0.80), followed by marmoset (79%). In rat, CHA containment (28% of volume ≥ 0.80) was comparable to an independent inter-atlas baseline (WHS regions within Duke Wistar parcels: 31%), demonstrating that boundary correspondence in rat is set by the available atlas infrastructure rather than by CHA (see Section 3.6 for more discussion). Mouse containment was perfect by construction. Together, the Dice and containment analyses establish that CHA respects established boundaries across species to the degree supported by existing atlas resources.

The biological validity of CHA’s regional definitions is supported by convergent evidence across independent modalities and species. Cross-species comparison of invasive tracer connectivity, using mouse anterograde and marmoset retrograde data mapped to CHA, revealed significant rank-order correspondence (Spearman ρ = 0.61, p < 10⁻¹¹), with agreement strengthening where marmoset tracer coverage was most complete (Fig. 6). A more extensive evaluation using diffusion MRI across all four species^31^ identified both conserved pathways, such as the entorhinal-hippocampal projection ranking as the most efficient in all species alongside thalamo-occipital and precentral-premotor connections, and lineage-specific divergence concentrated in anterior insula (human peak) and olfactory/piriform cortex (marmoset peak). Notably, the regions showing the greatest divergence in the diffusion MRI analysis are precisely those excluded from the tracer comparison due to absent marmoset injection coverage, making the two analyses complementary. These findings align with a broader pattern emerging across modalities: transcriptomic analyses report greater mouse-human similarity in sensorimotor than association cortex^32^, functional connectivity comparisons between human and macaque identify sensorimotor conservation alongside association divergence^33^, and white matter connectivity blueprints across primates reach similar conclusions^34,35^. That invasive tracer connectivity, transcriptomic correspondence, non-invasive imaging, and directed connectomics cost efficiency analysis independently converge on the same organizational principle, sensorimotor conservation paired with association divergence, provides convergent support that CHA captures biologically meaningful divisions.

The per-region homology confidence index (Fig. 7b) formalizes this assessment, with combined scores ranging from 0.82 to 0.98. The highest confidence regions (>0.95) are those with consistent nomenclature across all source atlases and strong geometric and connectivity correspondence, while the lowest scores identify where future data would be most informative. Regions relying on functional analogy rather than strict anatomical homology in mouse, particularly TEM_Inferior (ectorhinal cortex) and TEM_Superior (auditory cortex), received the lowest confidence scores (0.82), and geometric consistency analysis identified specific region pairs where cross-species spatial relationships diverge due to cortical folding differences, such as CIN_Posterior and PAR (R = 0.36). Connectivity-based evaluation could not be performed for anterior and posterior insula, anterior olfactory, piriform, or subcortical regions due to the cortical scope and incomplete coverage of the marmoset tracer dataset. These gaps define clear priorities for future work: primate tracing experiments targeting insular and olfactory cortex, and extension of connectivity validation to subcortical systems, would most improve confidence in the regions where it is currently lowest.

### 3.4. Utility of the common atlas framework

A central challenge in translational neuroscience is the absence of a standardized anatomical framework for relating findings across species. Species-specific atlases achieve high resolution within a single species but cannot, by definition, be directly compared across species. As a result, translational inferences from animal models to human neuroscience typically rely on informal reasoning about regional correspondence, with no systematic way to evaluate where such correspondence holds and where it breaks down. This gap is becoming more consequential: major international efforts are now extending invasive connectomics to primates, with systematic anterograde and retrograde tracing programs underway in marmoset through the Brain/MINDS project^37^ and related initiatives, representing a significant new commitment of resources and animal lives under an explicitly translational justification. Without a common anatomical framework, the field risks generating invaluable species-specific datasets that cannot be formally connected to one another or to the human brain they are ultimately intended to inform.

CHA addresses this gap by providing a unified coordinate system in which cross-species comparisons can be conducted quantitatively. The tracer validation presented here (Fig. 6) illustrates this directly: by mapping both mouse anterograde and marmoset retrograde tracer data to CHA, we identified where connectivity architecture is conserved across these species and where it diverges. This has practical consequences for experiment design. Where mouse-marmoset correspondence is strong, as in sensorimotor systems, primate-specific tracing experiments may not need to be repeated; where they diverge, as in association cortex, CHA pinpoints the regions most in need of targeted primate data. This is particularly relevant for subcortical systems: existing primate tracer resources, such as the Marmoset Cortico-Cortical Brain Connectivity^30^, contain labeled neurons in subcortical structures including thalamus but lack a standardized cross-species framework for assigning them. CHA bridges this gap by extending well-established mouse subcortical parcellation into primate space.

More broadly, CHA enables analyses that require one-to-one regional correspondence across species. A follow-up study^31^ combined CHA with mouse tracer-derived projection polarity and species-specific diffusion MRI tractography to construct directed connectomes across mouse, marmoset, rhesus macaque, and human. In this framework, CHA serves as the anatomical scaffold through which empirically validated directionality from mouse viral tracing is transferred to species where invasive data are unavailable, while each species’ own white-matter organization shapes the resulting efficiency landscape. This revealed conserved memory circuits alongside lineage-specific reallocations in temporal-insula-frontal, ventral visual stream, and olfactory pathways. Such directed cross-species pathway analyses depend on a common parcellation framework and would not be feasible with species-specific atlases alone.

Current deep learning approaches to brain segmentation are trained and validated within single species and cannot produce cross-species homologous parcellations across evolutionarily distant species because no training data for such mappings has existed. CHA provides the first standardized ground truth that could support training of cross-species segmentation models. With CHA labels defined consistently across mouse, marmoset, rhesus macaque, and human, it becomes feasible to train models that learn species-invariant anatomical features and generalize parcellation across species, potentially extending the framework to new species or imaging modalities without requiring manual delineation for each.

Similarly, recent transcriptomic efforts such as single-cell RNA sequencing have begun to characterize cell-type composition across species, revealing both conserved and divergent molecular signatures in homologous brain regions^32^. However, integrating these molecular data with macroscale anatomical organization requires a spatial framework that defines regional correspondence across species. CHA provides this framework: transcriptomic profiles can be mapped onto CHA regions to systematically compare cell-type composition across species within homologous divisions, linking molecular-level divergence to the macroscale organizational differences captured by volumetric, connectivity, and directed pathway analyses.

Together, these applications demonstrate that CHA serves not only as a descriptive resource but as an enabling infrastructure for quantitative cross-species neuroscience, providing a translational scaffold that connects invasive connectomics, transcriptomic mapping, and computational modeling across species to the human brain they are ultimately intended to inform.

### 3.5. Relationship to existing atlases

Many existing atlases have been developed independently within single species and use heterogeneous template-construction methods, making it difficult to achieve consistent anatomical definitions across resources. Although population-based templates such as ICBM2009a exist for humans^9^, and community templates are available for mouse and nonhuman primates, these resources were not designed for cross-species comparison and employ different processing pipelines, resolution standards, and nomenclature conventions. In addition, several cross-species and primate atlases have focused primarily on cortical surfaces or macro-architectural cortical divisions, with comparatively less standardized coverage of subcortical and deep gray structures essential for whole-brain analyses^5,7^. These differences in template construction and anatomical coverage contribute to the difficulty of establishing fully consistent whole-brain parcellations across species.

To ensure comparability across species, templates were selected or constructed according to consistent criteria: population-averaged minimal deformation construction (excluding single-subject or simple-average templates), high-resolution isotropic acquisition, standardized orientation with the origin at the anterior commissure, and open-access licensing. Where existing community templates met these criteria, they were adopted directly (ICBM2009a for human, Brain/MINDS MBM v3 with AC-centered reorientation for marmoset). Where they did not, new MDTs were constructed: For rhesus macaque, the NIMH Macaque Template (NMT v2)^38,39^ provides a well-constructed population average from 31 specimens, but its native coordinate system places the origin far from the anterior commissure and does not follow AC-PC alignment, requiring substantial spatial reorientation for cross-species consistency. We therefore constructed a new rhesus MDT from PRIMatE Data Exchange scans with the origin at AC and AC-PC alignment applied during template construction. For rodents, existing templates often retain non-standardized head orientations that complicate cross-species spatial correspondence. To facilitate adoption, we have warped widely used species-specific atlases into our MDT coordinate spaces and share them alongside the CHA labels.

It is worth noting that CHA does not replace existing species-specific resources; rather, it provides the missing translational layer that links them. Species-specific atlases remain the appropriate tool for within-species analyses at fine resolution. CHA operates at a coarser level where cross-species homology can be reasonably established, offering a coordinate system through which findings from one species can be formally related to another. By harmonizing parcellations across multiple species within a single framework, applying uniform tissue segmentation, and validating regional definitions against independent connectivity data, CHA complements the precision of species-specific atlases with the cross-species consistency that translational neuroscience requires.

### 3.6. Limitations and future directions

Several limitations of the current work should be acknowledged. Some structures, including the claustrum, substantia nigra, and subthalamic nucleus, were excluded because reliable segmentation was not feasible across MRI datasets for all species. The claustrum forms a thin, sheet-like structure between the insula and putamen, and both the substantia nigra and subthalamic nucleus are small deep nuclei with low intrinsic MRI contrast; even in humans, robust segmentation typically requires ultra-high-resolution or specialized contrast and extensive manual delineation^40–42^. Consequently, many widely used MRI-based atlases omit these nuclei or include them only when high-resolution data are available for a particular species. In the striatum, caudate-putamen separation proved difficult to identify in mouse MRI, requiring consolidation into a single caudoputamen parcel due to the absence of a clear anatomical boundary; the Allen Mouse Brain Atlas likewise treats caudoputamen as a single complex in its reference annotation^43^.

The mouse PAR parcel corresponds primarily to its continuous somatosensory territory, whereas primates possess a subdivided parietal cortex comprising postcentral, superior, inferior, and precuneus regions^33,44^. To preserve lobe-level comparability while acknowledging these anatomical differences, we retained a single undivided PAR parcel in mouse, designated as a parietal territory rather than lobe. While mouse somatosensory cortex corresponds to primate postcentral cortex, primate parietal association areas have no clear mouse homologue. Retaining PAR under a common label enables quantitative comparison: the tracer validation (Figure 6) revealed that sensorimotor connections to PAR are among the best-conserved between mouse and marmoset, while association connections to PAR are among the most divergent, a pattern that would be invisible if the region were excluded. To minimize nomenclature confusion, we have added parenthetical species-specific identifiers to mouse atlas labels where CHA names differ from standard mouse terminology (e.g., PAR(SSp/SSs), TEM_Superior(AUD)). These exclusions and consolidations reflect a combination of practical segmentation limitations and genuine differences in anatomical organization across species.

The rat is included as a core species in the framework, with a population-averaged MDT constructed from 18 Wistar specimens using the SAMBA pipeline, representing the largest such template reported for this species. Containment validation was performed against the Waxholm Space atlas^27,45^ (WHS; 133 cortical regions), which is based on a single Sprague Dawley specimen. The strain difference between Sprague Dawley (WHS) and Wistar (our MDT) may contribute to boundary misalignment. CHA containment of WHS regions (28% of volume ≥ 0.80) was comparable to an independent inter-atlas baseline computed by evaluating WHS containment within the Duke Wistar atlas^28^ (54 cortical regions), which yielded 31% of volume ≥ 0.80 (Figure 5d). This indicates that boundary correspondence is constrained by the available rat atlas infrastructure rather than by CHA itself. For mouse, the Allen Brain Atlas provides a comprehensive, community-standard whole-brain parcellation against which any parcellation can be benchmarked; no comparable resource currently exists for rat. Independent rat-specific parcellation validated against improved future atlas resources would strengthen confidence in the rat CHA.

Looking ahead, the atlas can be refined to capture finer levels of granularity. A promising next step is the introduction of Level 3 subdivisions in systems with pronounced functional specialization, such as the hippocampus, thalamus, and amygdala. Higher-resolution parcellations in these domains will enable the direct comparison of questions related to intra-system organization, such as hippocampal subfields or thalamic nuclei specializations, across species. Additionally, the mouse cohort used for MDT construction were male-only, which may influence volumetric scaling; extending the framework to include both sexes and across developmental stages is an important direction for future work. Such expansions will enhance the translational utility of the atlas, linking microcircuit-level distinctions to macroscale pathways and enabling more mechanistic models of brain organization.

### 3.7. Conclusion

In sum, the common hierarchical atlas offers a broadly applicable resource for comparative and translational neuroscience. It delivers consistent parcellation across divergent species, validated by independent invasive tracer data and supported by convergent evidence across modalities. By providing the missing cross-species coordinate system, CHA enables principled integration of diverse data types and formal evaluation of translational assumptions that have until now remained implicit.

## 4. Online Methods

### 4.1. Construction of Minimal Deformation Templates

To ensure anatomically consistent cross-species alignment, we constructed species-specific minimal deformation templates (MDTs). Except for humans (standard template), each MDT was generated separately, aligned to the AC–PC axis, and rigidly positioned with the origin at the anterior commissure (AC). This procedure follows the general principles of population-based template building implemented in ANTs, where iterative symmetric diffeomorphic registration is used to construct an unbiased minimal-deformation group average^11,12^. All atlas labels and analyses refer to these MDT spaces.

#### 4.1.1. Mouse MDT

All mouse procedures were approved by the Duke Institutional Animal Care and Use Committee (Duke IACUC approval A169-20-08). Male C57BL/6J mice were obtained from Jackson Laboratories and actively stained using a protocol involving transcardial infusion of ProHance in 10% buffered formalin, followed by post-fixation and equilibration^46,47^.This reduces T1 to ∼100 ms, enabling shorter TR and improved signal-to-noise ratio.

Imaging was performed on a 9.4 T vertical Oxford magnet with Resonance Research gradients (BFG 88/41) using a 13 × 25 mm solenoid coil tuned to 300 MHz. A Stejskal–Tanner sequence with compressed sensing achieved 25 µm isotropic resolution. We acquired 61 diffusion-weighted volumes (b = 3000 s/mm²) and 6 baselines (TR = 80 ms, TE = 14 ms) with uniformly distributed gradient directions. Compressed sensing^48,49^ (BART^50^) at 8× acceleration reduced the total acquisition time to ∼19 h per 4D dataset.

After acquisition, DWIs were corrected for eddy currents by registering each diffusion-weighted volume to the averaged baseline. Denoising used a local spatiotemporal noise model across k-and q-space^46^. Datasets were then reconstructed in DSI Studio using GQI.

The mouse MDT was constructed from 22 specimens (17 at 25 µm; 5 originally at 15 µm and downsampled to 25 µm). After preprocessing and coarse spatial checks, an initial template was formed via pairwise affine registrations: for each specimen, the set of affines to all others was averaged to obtain a median-space transform. Applying these transforms produced the initial mean. We then ran six iterative register-to-template cycles: (i) diffeomorphic registration to the current MDT, (ii) warping and averaging, (iii) averaging all deformation fields to visualize variability, and (iv) warp-dependent blurring proportional to mean deformation magnitude to focus updates on high-variance loci. This yielded a minimal-deformation population mean.

Labels in MDT space were generated by registering the final MDT to a symmetric mouse atlas and inverting the transforms. The final template was cropped and rigidly adjusted so the origin was at AC and AC/PC lay on a common axial slice.

#### 4.1.2. Rat MDT

Rat brain MRI were acquired at the Duke Center for In Vivo Microscopy (CIVM) under IACUC approval. Animals were actively stained using the same protocol as mice^46,47^. For rats, mandibles were removed to allow use of a smaller RF coil and improve sensitivity at high spatial resolution.

Specimens were scanned on a Bruker 7 T Avance III with high-performance gradients (Resonance Research, BFG 102/60; 1000 mT/m). Acquisition used a Stejskal–Tanner spin-echo sequence (TR = 100 ms, TE = 22.5 ms, b = 3000 s/mm²) with 61 diffusion directions uniformly distributed on the sphere; a b0 image was acquired after every 10th diffusion-weighted scan. Data were streamed to an HPC cluster and reconstructed with compressed sensing^48,49^. Baselines were averaged; all DWIs were then registered to the baseline with ANTs^46,51^.

A rat MDT was constructed from 18 Wistar rats (∼P90) using the SAMBA pipeline^52^, built on ANTs. Registration was based on DWI contrast. To obtain symmetric labels, the final MDT was registered to the symmetric Wistar rat atlas, and inverse transforms were applied to warp the label set into MDT space. The final MDT was cropped and rigidly adjusted so the origin was at AC and AC/PC lay on a common axial slice.

#### 4.1.3. Marmoset MDT

MRI data were obtained from the Brain/MINDS Marmoset Brain MRI dataset^53^. Marmoset atlas data were derived from Marmoset Brain Mapping (MBM) v3^10^. We applied minimal spatial header adjustments to achieve AC-centered coordinates. Specifically, the NIfTI srow matrix was modified to (units in mm):

0 0 -14.6
0.2 0 -25.7
0 0.2 -13.3

These changes preserve MBM v3 image content while placing the AC at (0,0,0) and maintaining isotropic 0.2 mm voxels. Subsequent rigid checks ensured AC–PC alignment.

#### 4.1.4. Rhesus Macaque MDT

Rhesus macaque T1-weighted images were derived from The PRIMatE Data Exchange (PRIME-DE) via International Neuroimaging Data-sharing Initiative (INDI)^54^. Each volume was rigidly aligned to AC–PC, with the origin set at AC. We constructed a minimal deformation template (MDT) using an iterative diffeomorphic averaging procedure implemented in ANTs^12^. The final template was recentered at the anterior commissure (AC) and resampled to isotropic resolution to facilitate label transfer.

#### 4.1.5. Human MDT

For humans, we adopted the widely used ICBM152 Nonlinear Asymmetrical 2009a template (ICBM2009a)^9^ as the species reference space. This template is a population-average of 152 adult (age 18-44y) T1-weighted MRI volumes, generated using iterative nonlinear registration to minimize anatomical variability across individuals. The final template was AC–PC aligned, with the origin set at the anterior commissure (AC), and provides an established standard for human neuroimaging studies.

### 4.2. Construction of Brain parcellations

Tissue (structural compartment) segmentation was performed in U-NET Studio^55^ to generate probability maps for gray matter, white matter, deep nuclei, cerebellum, and CSF from high-resolution MRI. The common grey matter parcellation framework was subsequently generated in DSI Studio^56^. Structural compartment segmentation was performed using a cascaded cross-species training procedure, with separate hierarchies for primate and rodent lineages. Within each lineage, the species with the largest brain (and correspondingly superior signal quality) served as the initial model. For primates, the human-trained model was first applied to segment the rhesus macaque template; after manual inspection and correction, the refined rhesus segmentations were used to train the marmoset model. For rodents, the rat-trained model was applied to segment the mouse template, with manual correction before finalizing the mouse model. This leader-follower approach leverages the higher signal quality of larger brains to bootstrap segmentation in smaller species while ensuring anatomical accuracy through manual verification at each step.

Subsequently, a two-level hierarchy of common regions of interest across species was defined for the grey matter tissue compartment and deep grey matter structures. Level-1 parcellation comprised nine cortical and subcortical divisions, delineated using major anatomical boundaries such as cortical lobes and clearly identifiable deep structures in high-resolution MRI. The nine Level-1 macro-regions were delineated using major anatomical landmarks identifiable in high-resolution MRI. In humans, the central sulcus defined the frontal-parietal boundary, the Sylvian (lateral) fissure separated temporal cortex and exposed the insula, the parieto-occipital sulcus marked the parietal-occipital boundary, and the cingulate sulcus delimited the medial extent of cingulate cortex. Basal ganglia and thalamic divisions followed clearly identifiable nuclear margins. In nonhuman primates, homologous sulcal landmarks were used where present (central sulcus, lateral sulcus, parieto-occipital sulcus), supplemented by atlas-defined boundaries from CIVM (rhesus) and Paxinos/MBM (marmoset). In rodents, where cortical sulci are largely absent, Level-1 boundaries were established by registering the Allen Brain Atlas to the mouse MDT and grouping ABA regions into the nine macro-divisions based on literature-guided functional and positional correspondence. The rat Level-1 parcellation was derived by warping the mouse CHA to the rat MDT.

Level-2 further subdivided Level-1 regions along secondary anatomical margins such as cortical gyri, sulci, or known cytoarchitectonic transitions. Subdivisions were primarily established in humans using high-resolution MRI and aligned with anatomical conventions from widely used atlases. Homologous regions in animals were then identified using a literature-guided approach, incorporating prior anatomical and histological studies, species-specific atlases, and patterns of regional naming and usage in the comparative neuroimaging literature.

Because cross-species homology is not always uniquely defined, particularly in higher-order association areas, the framework prioritized three principles: (i) preserve regional definitions widely used in existing atlases, (ii) maximize cross-species consistency where boundaries could be reasonably identified in MRI, and (iii) retain functional interpretability. In some cases, functional considerations were decisive. For example, the superior temporal parcel combines the superior and middle temporal gyri in humans, which cannot be reliably separated in nonhuman primates or rodents but together form a coherent auditory–language system. By contrast, the superior, middle, and inferior frontal gyri, though clearly identifiable in humans, were not retained as separate parcels due to the absence of clear homologues in lower species. Instead, functionally grounded regions such as precentral, premotor, and postcentral were emphasized to preserve sensorimotor comparability. Similarly, olfactory parcels were included despite their small size in humans, reflecting their critical role in rodent and marmoset cognition and a rich comparative literature. The medial temporal parcel in mouse encompasses the rhinal complex (entorhinal, perirhinal, postrhinal), while in primates it includes the entorhinal and parahippocampal cortices; across species, this region mediates hippocampal–neocortical interactions. Conversely, structures such as the claustrum, substantia nigra, and subthalamus, while homologous in principle, were excluded due to segmentation challenges in cross-species MRI datasets.

Primary species (human, rhesus macaque, marmoset, and mouse) were delineated directly from MRI-based parcellations. To extend coverage, rat parcellations were derived by warping the mouse atlas, using MRI-based alignment. These warped parcellations provided coverage for additional species while preserving consistency with the core cross-species framework.

For each Level-2 region, homologues were identified from one or more standard atlases: FreeSurferSeg (human)^57,58^, Brodmann (human)^59^, the Allen Brain Atlas (mouse)^43^, CIVM rhesus atlas (rhesus macaque)^60^, Inia19–NeuroMaps (rhesus macaque)^61^, Brain/MINDS (marmoset)^62^, MBM atlas (marmoset)^63^, and Paxinos (marmoset)^64^, with priority given to sources providing the clearest anatomical or functional correspondence. Where multiple atlases were necessary (e.g., in marmoset and rhesus), boundaries were cross-validated across references. Detailed listings of atlas sources, supporting references, and region coverage for each Level-2 parcel are provided in Table 1.

### 4.3. Validation of the Common Atlas Across Scales and Species

#### 4.3.1. Validation of Human Parcellation Using Neuroparc Atlases

To evaluate the anatomical consistency of the common human atlas, we compared each Level-2 region against corresponding regions from standardized human parcellations in the Neuroparc repository^4^. Both CHA and the Neuroparc atlases are defined in ICBM152 standard space^9^, so no inter-atlas registration was required. The CHA human atlas (0.5 mm isotropic resolution) was downsampled to 1 mm isotropic to match the highest available Neuroparc resolution prior to comparison. Candidate matches were identified by flexible text-string matching of region labels and cross-checked against anatomical conventions. Binary masks were constructed for each atlas, and Dice similarity coefficients were computed to quantify voxelwise overlap. Neuroparc atlases with poor performance for each region, defined as an average Dice overlap < 0.1 across other Neuroparc atlases, were excluded from region-level comparisons.

For each common region, Dice distributions were summarized as "Common vs. Others" (overlap between CHA and each Neuroparc atlas) and "Others vs. Others" (overlap among the Neuroparc atlases themselves, excluding CHA). Region-level summary metrics included the median Dice similarity for each comparison type and the difference between them (ΔMedian).

To test whether CHA achieves at least equivalent spatial overlap with established atlases as those atlases achieve with each other, we computed the per-region ΔMedian Dice (median of "Common vs. Others" minus median of "Others vs. Others") and applied a one-sided Wilcoxon signed-rank test across regions, testing the alternative hypothesis that ΔMedian ≥ 0. We additionally performed a non-inferiority test by shifting the ΔMedian values by pre-specified margins (0.05 and 0.10) and testing whether the shifted values remained significantly above zero. This non-inferiority framing reflects the fact that no single parcellation can serve as ground truth for cortical boundary definitions given the well-documented inconsistencies across established human brain atlases^1,3,4^.

#### 4.3.2. Evaluation Against Mouse Allen Brain Atlas Major Regions

To evaluate correspondence between the common mouse parcellation and its source atlas, we compared Level-2 regions against the 13 major divisions of the Allen Brain Atlas (ABA). Dice similarity coefficients were computed between each ABA major parcel and the set of common ROIs. Because CHA mouse regions were constructed within the same MDT space to which the ABA was registered, no additional alignment was required. High Dice values indicate direct one-to-one correspondence between ABA and CHA regions (e.g., thalamus, hypothalamus, hippocampus, olfactory areas). Lower Dice values for broad ABA parcels such as isocortex reflect the intentional subdivision of mouse cortical territory into human-centric divisions to enable cross-species comparison, rather than boundary misalignment. Statistical comparison of CHA against ABA is not performed, as the two atlases serve different purposes: ABA provides a comprehensive mouse-specific parcellation, while CHA reorganizes mouse cortical territory into a framework directly comparable with primates and humans.

#### 4.3.3. Containment Validation Using Finer-Scale Species-Specific Atlases

Containment scores were used to test whether the coarser common atlas respected the boundaries of finer-scale species-specific parcellations. For each species, the relevant species-specific atlas was registered to the species MDT template space using nonlinear transformations prior to CHA construction. Because CHA was subsequently built within these same MDT spaces, the resulting CHA labels and the warped species-specific labels share the same coordinate system, and no additional alignment was required for validation.

Validation was performed for mouse against the ABA (372 regions), for rat against the Waxholm Space atlas (WHS v4.01; 133 cortical gray matter regions)^27^, for marmoset against the MBM atlas (122 regions), and for rhesus against the CIVM atlas (220 regions). For rat, we additionally computed an independent inter-atlas baseline by evaluating WHS containment within the Duke CIVM Wistar atlas (54 regions)^28^, providing a reference for inter-atlas agreement independent of CHA. Note that the WHS atlas is based on a single Sprague Dawley specimen, whereas both the CHA rat MDT (18 Wistar specimens) and the Duke atlas are Wistar-derived; the resulting strain difference may contribute to boundary misalignment between WHS and either Wistar-based atlas.

For each finer ROI, we computed the voxelwise fractional overlap with all target parcels and retained the maximum containment score, ensuring that each ROI was matched to its best-fitting target label while ignoring partial overlaps with non-dominant regions. Because target atlases differ in the number and size of parcels, larger parcels have higher chance containment. To correct for this, we applied a specificity adjustment analogous in structure to the adjusted Rand index^65^ used in cluster analysis: *adjusted* = (*raw* − *expected*) / (1 − *expected*), where *raw* is the maximum fractional overlap and *expected* is the volume fraction of the best-matching target parcel relative to total labeled gray matter volume. This yields a score of 1 for perfect containment, 0 for containment no better than chance given parcel size. All reported containment values and ECDF curves use this adjusted metric, weighted by finer-atlas region volume.

Additionally, to test anatomical resilience under spatial perturbation, common labels were rigidly rotated about a central axis in fixed angular increments, without resampling or interpolation. Containment scores were recomputed for each rotated configuration using the same procedure, enabling direct comparison of distributional shifts across angles.

#### 4.3.4. Cross-Species Connectivity Validation

To validate CHA regional definitions using independent connectivity data, we compared directed tracer connectivity matrices derived from mouse and marmoset. Mouse connectivity was computed from ∼1,197 anterograde viral tracing experiments from the Allen Mouse Brain Connectivity Atlas^29^, mapped to CHA as described in^31^. Marmoset connectivity was derived from 143 retrograde fluorescent tracer injections from the Marmoset Brain Connectivity Atlas^30^, accessed via the public API (http://www.marmosetbrain.org). For each injection, the fraction of extrinsic labeled neurons (FLNe) was obtained for all source areas, quantifying the strength of projections from each cortical area to the injection site.

The marmoset FLNe matrix, originally defined in Paxinos stereotaxic space^22^, was converted to CHA space using containment-weighted transformation. For each Paxinos area, we computed the fractional overlap with each CHA region from the MBM_vPaxinos atlas registered to the marmoset MDT. The CHA-space matrix was then computed as ***F***_*CHA*_ = ***W***^*T*^ ***F* *W***, where ***F*** is the Paxinos-space FLNe matrix and ***W*** is the Paxinos-to-CHA containment fraction matrix. Label mismatches between the API output and the atlas were resolved through manual name matching (e.g., "A1/2" to "A1/A2", "V5(MT)" to "V5").

The comparison was restricted to 11 cortical CHA regions with adequate marmoset tracer coverage. Four cortical regions (INS_Anterior, INS_Posterior, OLF_Anterior, OLF_Piriform) were excluded because they lacked systematic injection coverage in the marmoset dataset (zero or near-zero inflow columns), meaning any signal in those regions reflected incidental labeling rather than quantified connectivity. Subcortical CHA regions were excluded because the marmoset tracer dataset is restricted to corticocortical connections. Only directed pairs where both matrices had nonzero entries were included in the analysis.

Cross-species correspondence was quantified using Spearman rank correlation between mouse and marmoset directed connection weights across all included pairs. To assess whether correspondence scaled with the quality of marmoset tracer coverage, directed pairs were divided into quartiles based on the minimum marmoset signal (total inflow plus outflow) of the two regions in each pair. Per-connection agreement was computed as 1 minus the absolute difference in rank percentiles between mouse and marmoset for each directed pair: 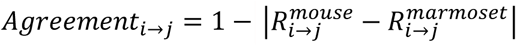, where 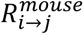 *a*nd 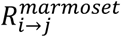 are the within-species rank percentiles of connection weight for the directed pair *i* → *j*. Values near 1 indicate similar relative ranking across species; values near 0 indicate divergent ranking. Per-region agreement was computed as the mean across all directed pairs involving that region. Sorted per-connection agreement values are provided in Supplementary Table 3. A more extensive non-invasive connectivity-based validation and comparison across all four species is presented in a follow-up study using species-specific diffusion MRI tractography^31^.

#### 4.3.5. Per-Region Homology Confidence Assessment

To quantify the confidence of cross-species regional assignments, we developed a composite homology confidence index integrating three independent criteria for each CHA region: nomenclature directness, geometric consistency, and cross-species connectivity correspondence.

##### Nomenclature directness

For each CHA region in each species, we scored the degree to which established species-specific atlases explicitly support the regional assignment on a 0–3 scale: 3 = the source atlas uses the same or directly equivalent term (e.g., "hippocampus" in all species); 2 = the atlas uses a recognized anatomical equivalent requiring minimal interpretation (e.g., Allen Brain Atlas MOp mapped to CHA FRO_Precentral via established primary motor correspondence); 1 = the mapping relies on a functional analog supported by published literature (e.g., mouse ectorhinal cortex mapped to CHA TEM_Inferior based on connectivity and functional response properties^13,14^); 0 = no direct atlas support. Scores were assigned based on the atlas sources documented in Table 1. The per-region nomenclature score was computed as the mean across species, normalized to a 0–1 scale.

##### Geometric consistency

For each species, we computed the centroid (center of mass in world coordinates) of each CHA region from the species’ atlas label volume. For every pair of CHA regions (*i*, *j*), we computed the three-dimensional direction vector *d*_*s*_ = *centroid*_*j*_ − *centroid*_*i*_ in each species *s*. Each direction vector was normalized to unit length 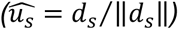 to remove the influence of brain size, so that each species contributes equally regardless of absolute inter-regional distances. The mean resultant length^36^ R was then computed as: 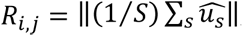, where *S* is the number of species, and *R* ranges from 0 to 1. When all species’ direction vectors point in the same direction, the unit vectors sum constructively and *R* approaches 1. When direction vectors are scattered across species, the unit vectors partially cancel and *R* approaches 0. The per-region geometric consistency score reported in Table 1 and Figure 7b was computed as the mean *R* across all region pairs involving that region.

##### Cross-species connectivity correspondence

Using the cross-species tracer comparison described above, we computed a per-region connectivity score as the mean rank-percentile agreement across all directed pairs involving that region for which both mouse and marmoset matrices had nonzero entries. Four cortical CHA regions lacking systematic marmoset injection coverage (INS_Anterior, INS_Posterior, OLF_Anterior, OLF_Piriform) and all subcortical regions (which fall outside the scope of the corticocortical marmoset tracer dataset) were marked as not evaluable for this criterion.

##### Combined index

The homology confidence index for each region was computed as the mean of all available criteria (each scaled 0–1). Regions for which a criterion could not be evaluated were scored on the basis of the remaining criteria and explicitly marked.

## Acknowledgments

Computational resources were provided by the Pittsburgh Supercomputing Center through NSF ACCESS allocations MED230052 and CIS200026.

## Use of AI Tools

During the preparation of this work the authors used generative AI tools to assist with manuscript drafting, literature review, and organizational structuring. After using these tools, the authors reviewed and edited the content as needed and take full responsibility for the content of the published article.

## Author Contributions

*Siva Venkadesh:* Conceptualization, Methodology (Level-1/2 parcellations, cross-species homology definitions), Formal analysis, Software, Validation, Visualization, Writing – Original Draft, Writing – Review & Editing, Supervision

*Yuhe Tian*: Software, Validation, Formal analysis, Visualization

*Wen-Jieh Linn*: Data Curation, Visualization

*Jessica Barrios Martinez*: Data Curation, Visualization, Writing – Review & Editing

*Harrison Mansour*: Data Curation, Resources, Writing – Review & Editing

*David J. Schaeffer*: Data Curation, Resources, Writing – Review & Editing

*Diego Szczupak*: Data Curation, Resources, Writing – Review & Editin

*Afonso C. Silva*: Methodology (marmoset), Resources, Writing – Review & Editing

*G Allan Johnson*: Conceptualization, Methodology (mouse, rat), Study oversight, Resources, Writing – Review & Editing, Funding acquisition.

*Fang-Cheng Yeh*: Conceptualization, Methodology (Level-0 tissue segmentation), Study oversight, Project administration, Resources, Writing – Review & Editing, Funding acquisition.

## Data Availability

The atlas data presented here are freely available at https://github.com/data-others/atlas for each species *(<species>_CHA.nii.gz)*

## Code Availability

All analysis and figure-generation code supporting the findings of this study are archived at Zenodo (https://doi.org/10.5281/zenodo.17653308) in the directory *projects/common_cross_species_atlas/*.

