## Supplementary Figures for "A hierarchical framework for cortical and subcortical gray-matter parcellation across rodents, primates, and humans"

| 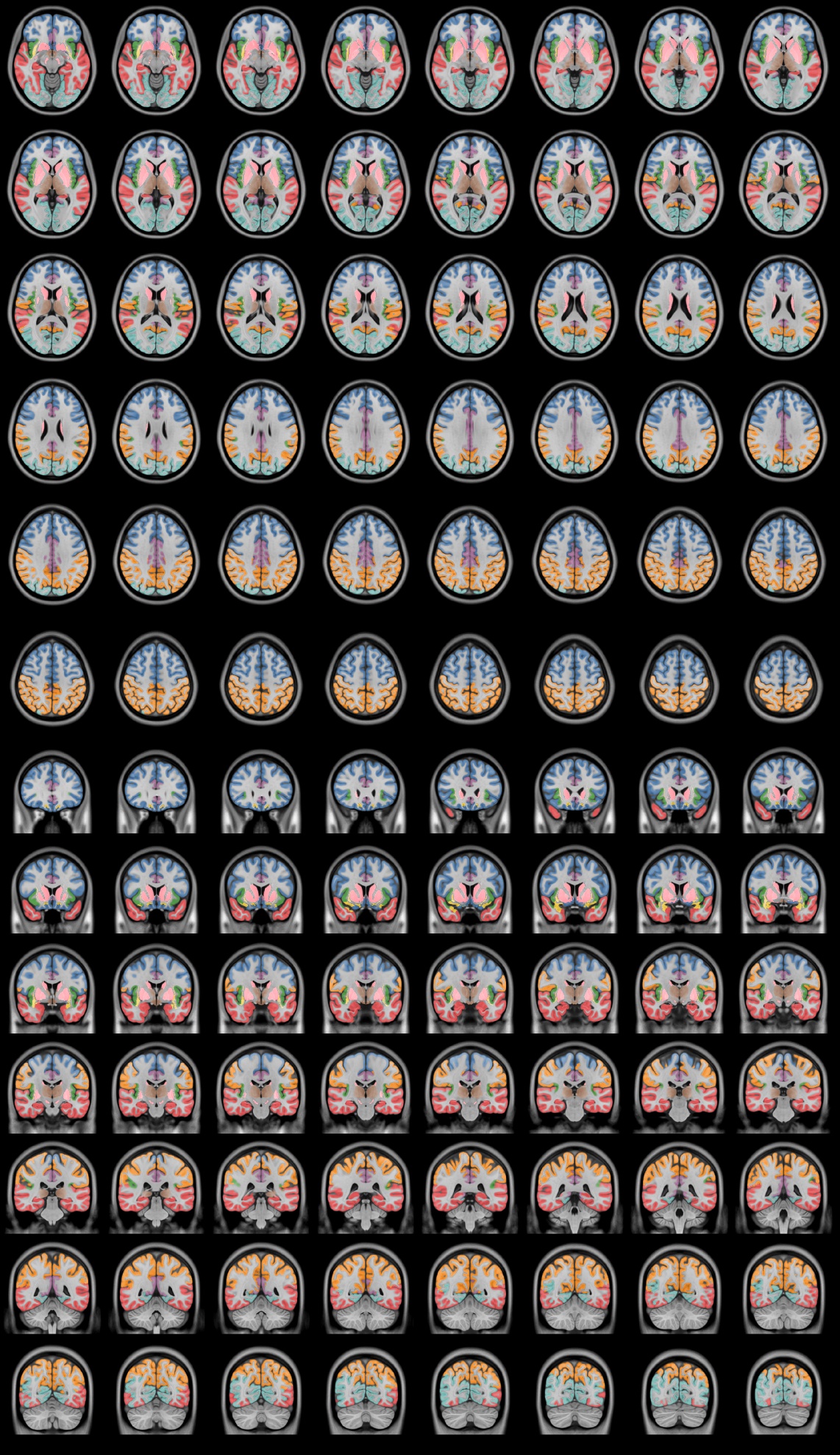 | **Supplementary Figure 1 \| Human atlas mosaic views.**  Axial and coronal mosaic views of the Level-1 cortical parcellation in the human brain. Each color corresponds to a macro-region as defined in the common cross-species atlas and shown in Figure 2 (frontal, parietal, temporal, occipital, insular, cingulate, basal ganglia, thalamic, and olfactory divisions). Renderings highlight the spatial extent and anatomical boundaries of these divisions in stereotaxic coordinates. |
| --- | --- |

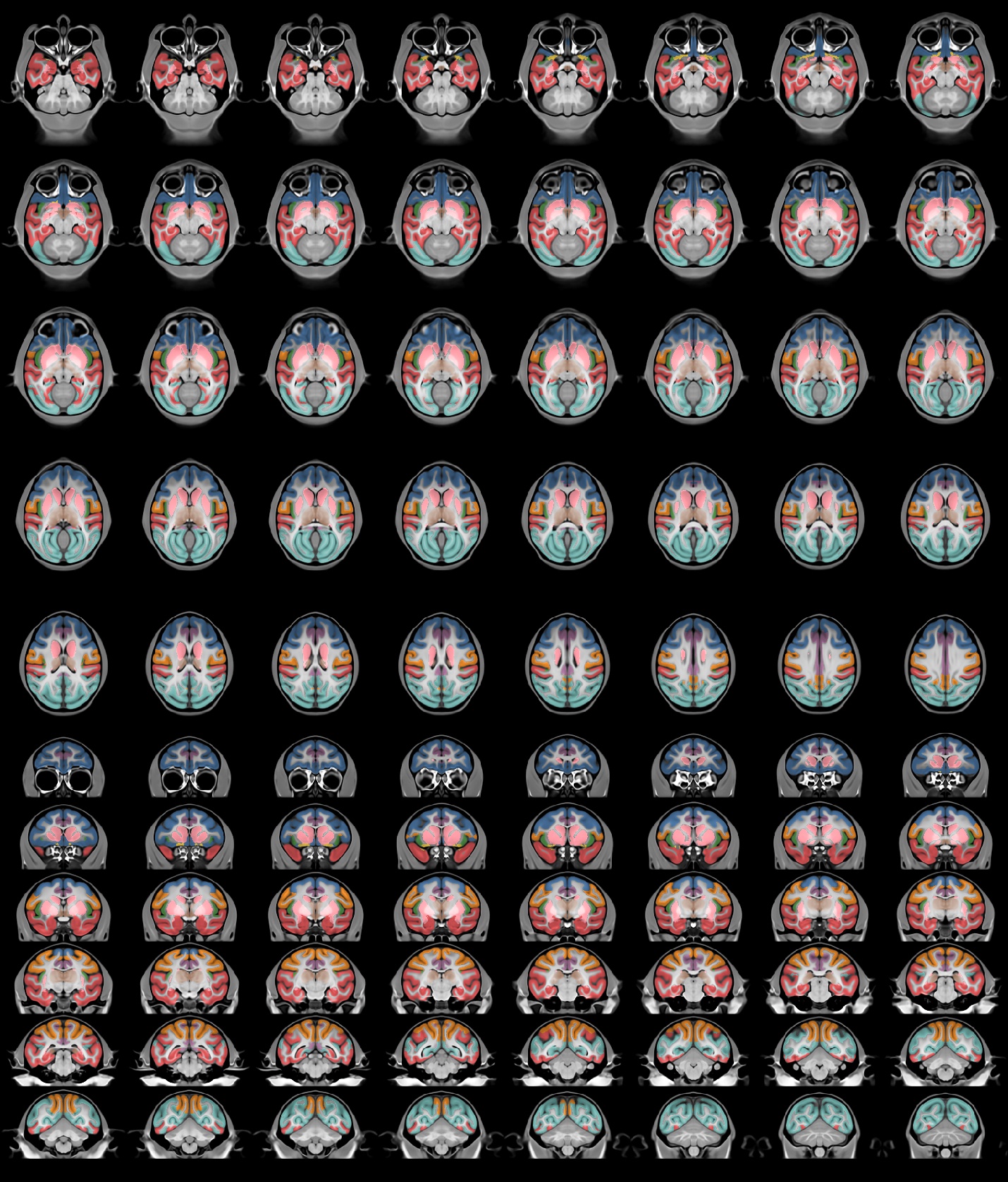

**Supplementary Figure 2 | Rhesus macaque atlas mosaic views.**

Axial and coronal views of the Level-1 parcellation in rhesus macaque, using the same color scheme as Figure 2. Mosaics illustrate overall alignment of macro-regions with human homologues while showing species-specific sulcal and gyral morphology.

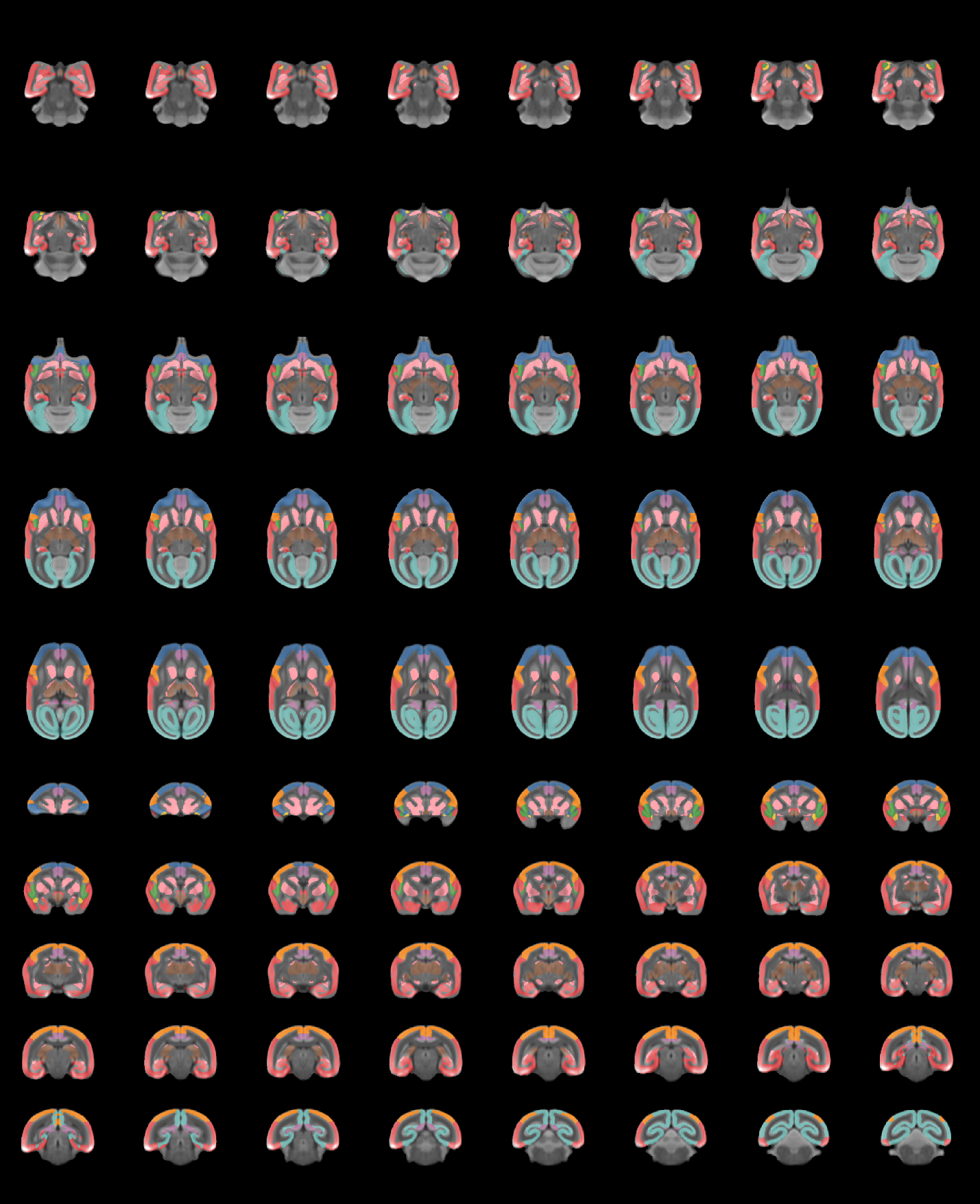

**Supplementary Figure 3 | Marmoset atlas mosaic views.**

Level-1 parcellation of the marmoset brain displayed in axial and coronal mosaics. Colors follow the common scheme in Figure 2, emphasizing conserved macro-organization and the more compact cortical layout characteristic of New World primates.

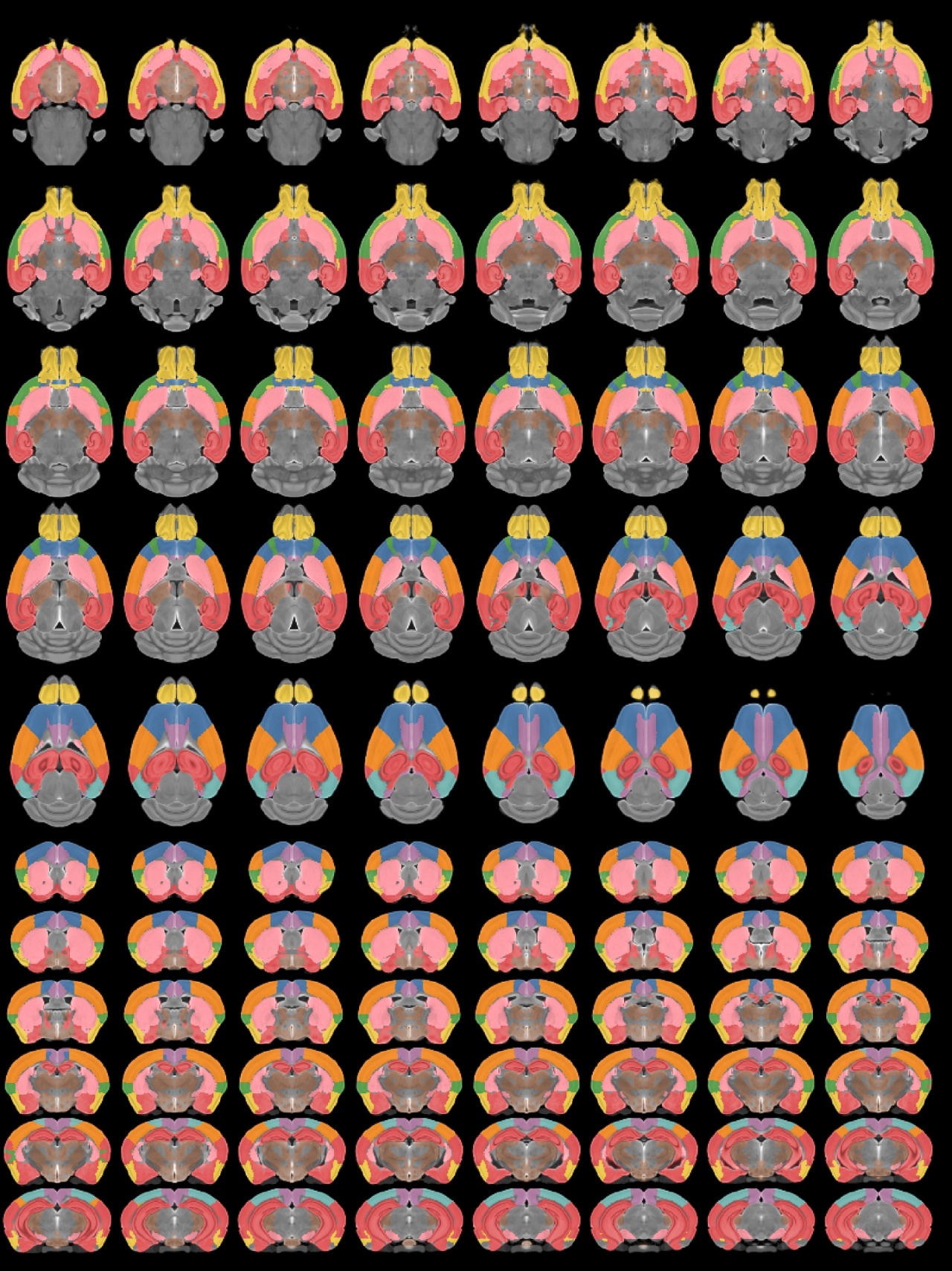

**Supplementary Figure 4 | Mouse atlas mosaic views.**

Axial and coronal mosaic views of the Level-1 parcellation in the mouse brain. Colors correspond to the same macro-regions used in Figure 2, demonstrating cross-species continuity of frontal, parietal, temporal, occipital, insular, cingulate, basal ganglia, thalamic, and olfactory divisions within the shared atlas framework.

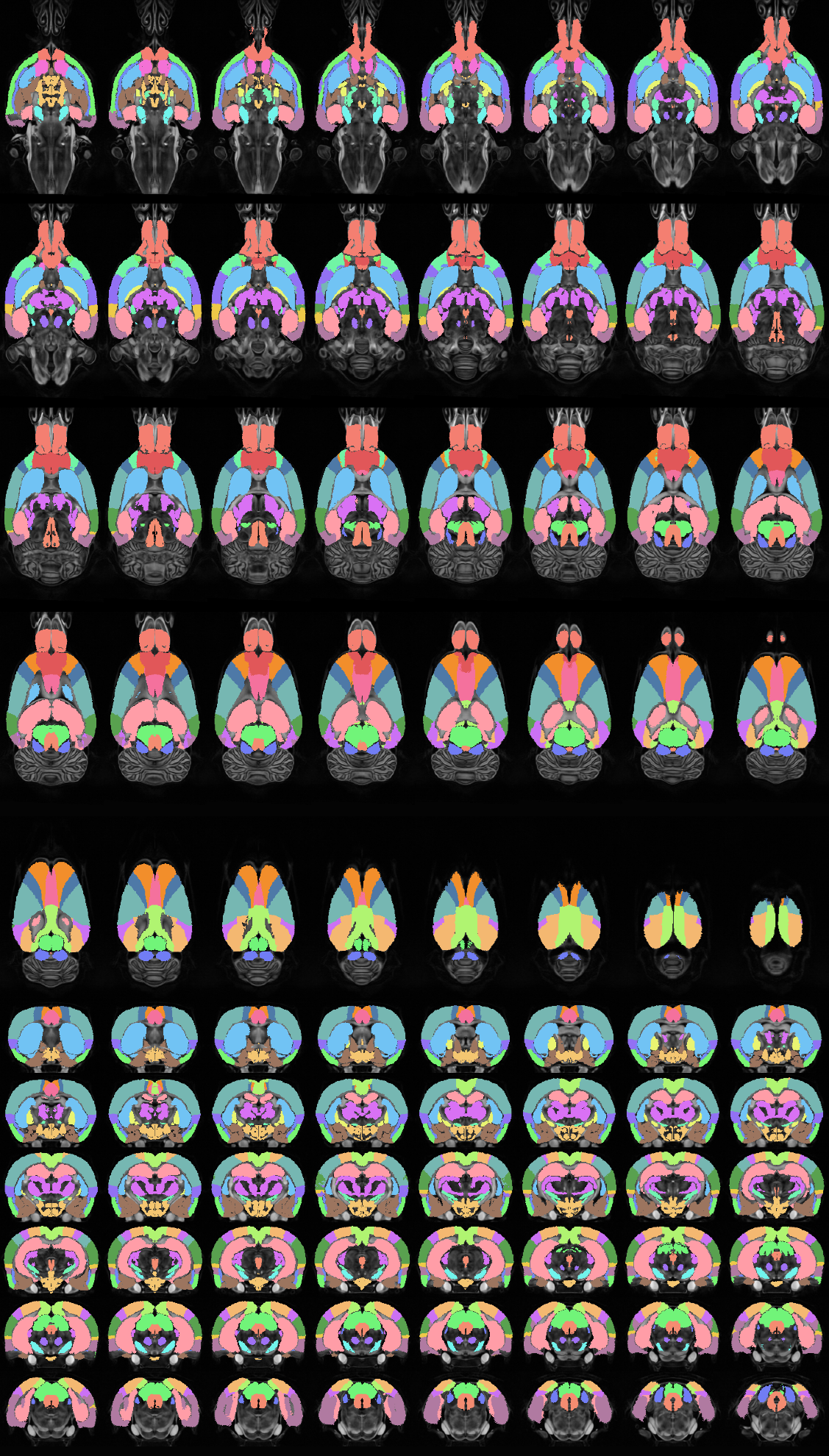
**Supplementary Figure 5 | Rat atlas mosaic views.**

Axial and coronal mosaic views of the Level-1 parcellation in the rat brain. Rat parcellations were derived by warping the mouse CHA using MRI-based nonlinear registration, as described in Methods.

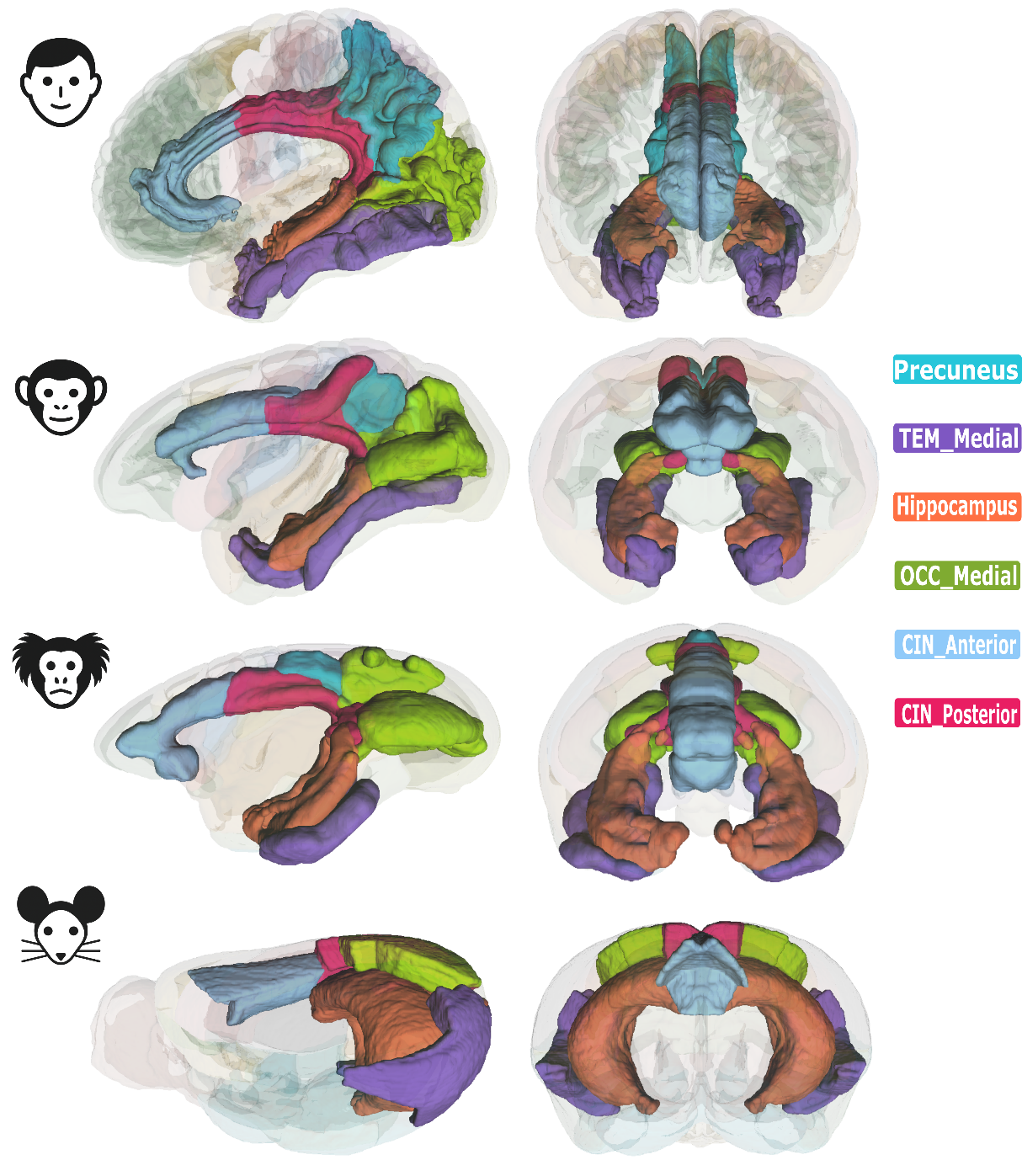

**Supplementary Figure 6 | Medial structures across species.** Three-dimensional renderings of medial CHA regions shown in sagittal (left) and coronal (right) views for human, rhesus macaque, marmoset, and mouse. Regions displayed: precuneus (cyan; primates only), medial temporal cortex (purple), hippocampus (orange), medial occipital cortex (green), anterior cingulate (light blue), and posterior cingulate (pink). The dorsomedial-to-ventromedial repositioning of posterior cingulate from rodent to primate is apparent in both views, consistent with known cortical folding differences and the low geometric consistency for the CIN_Posterior-PAR pair (R = 0.36; Figure 7a). Brain surfaces are rendered semi-transparently to reveal the depth of medial structures.

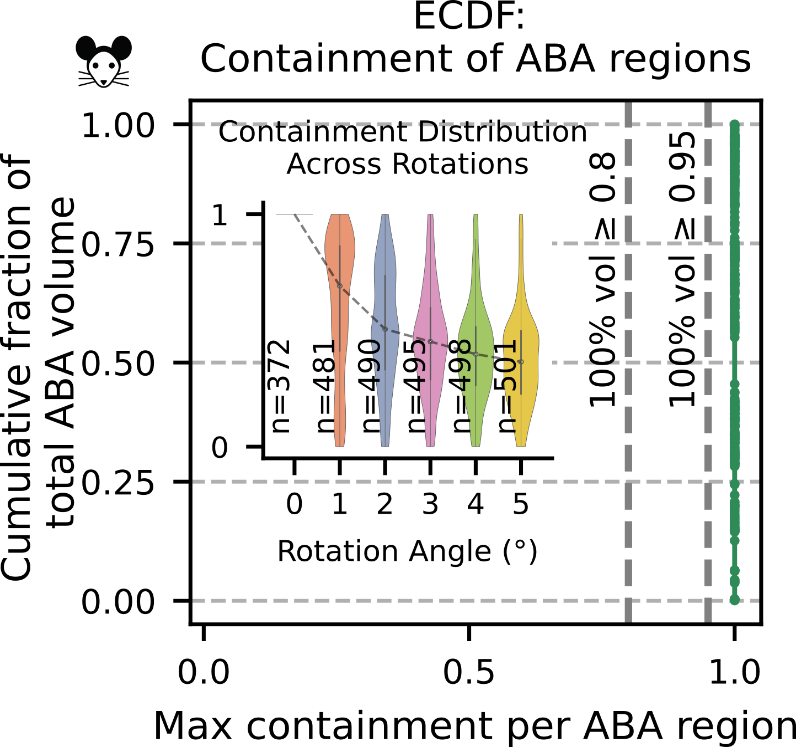

**Supplementary Figure 7 | Mouse containment validation.** Volume-weighted ECDF of containment scores for Allen Brain Atlas (ABA) regions within CHA. All regions achieved perfect containment (score = 1.0), as expected given that ABA served as the sole source for CHA mouse labels (no boundary loss). Rotation perturbation analysis (inset) demonstrates systematic decay of containment with increasing spatial disruption, validating the sensitivity of the metric.
