## Supplementary Table 1 for "A hierarchical framework for cortical and subcortical gray-matter parcellation across rodents, primates, and humans"

**Supplementary Table 1.** Abbreviations used in the manuscript, organized by category.

| **Abbreviation** | **Full Term** |
| --- | --- |
| ***Atlas sources*** | |
| ABA | Allen Brain Atlas (mouse) |
| BA | Brodmann areas (human) |
| BMA | Brain/MINDS Marmoset Reference Atlas |
| CIVM | Center for In Vivo Microscopy atlas (rhesus macaque) |
| FSS | FreeSurferSeg (human structural segmentation) |
| INM | INIA19–NeuroMaps atlas (rhesus macaque) |
| MBM | Marmoset Brain Mapping Atlas |
| Paxinos | Paxinos Histological Atlas (marmoset) |
| WHS | Waxholm Space atlas (rat) |
| ***Tissue classes (Level 0)*** | |
| GM | Gray Matter |
| WM | White Matter |
| DGM | Deep Gray Matter |
| CBL / CBGM | Cerebellum / Cerebellar Gray Matter |
| CSF | Cerebrospinal Fluid |
| ***Level-1 macro-regions*** | |
| FRO | Frontal Lobe |
| PAR | Parietal Lobe |
| TEM | Temporal Lobe |
| OCC | Occipital Lobe |
| INS | Insular Cortex |
| OLF | Olfactory Cortex |
| CIN | Cingulate Cortex |
| BG | Basal Ganglia |
| THL | Thalamic Areas |
| ***Level-2 subdivisions: Frontal*** | |
| PreC / FRO_Precentral | Precentral (primary motor) cortex |
| PreM / FRO_Premotor | Premotor (secondary motor) cortex |
| PrF / FRO_Prefrontal | Prefrontal cortex |
| ***Level-2 subdivisions: Parietal*** | |
| PAR | Parietal lobe (undivided in rodents) |
| PAR_Postcentral | Postcentral (primary somatosensory) cortex (primates) |
| PAR_Superior | Superior parietal lobule (primates) |
| PAR_Inferior | Inferior parietal lobule (primates) |
| PAR_Precuneus | Precuneus (primates) |
| ***Level-2 subdivisions: Temporal*** | |
| TemS / TEM_Superior | Superior temporal cortex |
| TemI / TEM_Inferior | Inferior temporal cortex |
| TemM / TEM_Medial | Medial temporal cortex (entorhinal/parahippocampal) |
| Hipp / TEM_Hippocampus | Hippocampus |
| Amyg / TEM_Amygdala | Amygdala |
| ***Level-2 subdivisions: Occipital*** | |
| OccL / OCC_Lateral | Lateral occipital cortex |
| OccM / OCC_Medial | Medial occipital cortex |
| ***Level-2 subdivisions: Insular*** | |
| InsA / INS_Anterior | Anterior insula |
| InsP / INS_Posterior | Posterior insula |
| ***Level-2 subdivisions: Olfactory*** | |
| OlfA / OLF_Anterior | Anterior olfactory cortex |
| Pir / OLF_Piriform | Piriform cortex |
| ***Level-2 subdivisions: Cingulate*** | |
| CingA / CIN_Anterior | Anterior cingulate cortex |
| CingP / CIN_Posterior | Posterior cingulate / retrosplenial cortex |
| ***Level-2 subdivisions: Basal Ganglia*** | |
| CPu / BG_CaudoPutamen | Caudoputamen (undivided in rodents; caudate + putamen in primates) |
| NAc / BG_Accumbens | Nucleus accumbens |
| Pall / BG_Pallidum | Globus pallidus |
| ***Level-2 subdivisions: Thalamic*** | |
| Thal / THL_Thalamus | Thalamus |
| Hypo / THL_Hypothalamus | Hypothalamus |
| ***Template and methodological terms*** | |
| MDT | Minimal Deformation Template |
| ICBM2009a | ICBM Nonlinear Asymmetric Human Template (2009a) |
| AC | Anterior commissure |
| AC–PC | Anterior commissure – Posterior commissure axis |
| CHA | Common Hierarchical Atlas |
| CDF | Cumulative Distribution Function |
| ROI | Region of Interest |
| U-Net | Convolutional neural network architecture used for tissue segmentation |
| SAMBA | Small Animal Multivariate Brain Analysis pipeline |
| ANTs | Advanced Normalization Tools |
| ***Imaging and acquisition terms*** | |
| dMRI | Diffusion magnetic resonance imaging |
| T1w / T2w | T1-weighted / T2-weighted MRI |
| GQI | Generalized q-sampling imaging |
| HARDI | High angular resolution diffusion imaging |
| DWI | Diffusion-weighted image |
| TR / TE | Repetition time / Echo time |
| ***Species abbreviations (used in Table 1)*** | |
| Mo | Mouse |
| Mz | Marmoset |
| Rh | Rhesus macaque |
| Hu / H | Human |
| ***Validation and statistical terms*** | |
| Dice | Dice similarity coefficient |
| R | Mean resultant length (geometric consistency) |
| FLNe | Fraction of extrinsic labeled neurons |
| Geo. R | Geometric consistency score (per-region mean of pairwise R) |
| Nom. | Nomenclature directness score |
